# The effect of depth variation on disparity tasks in natural scenes

**DOI:** 10.1101/162222

**Authors:** Arvind V. Iyer, Johannes Burge

## Abstract

Local depth variation is a distinctive property of natural scenes and its effects on perception have only recently begun to be investigated. Here, we demonstrate how natural depth variation impacts performance in two fundamental tasks related to stereopsis: half-occlusion detection and disparity detection. We report the results of a computational study that uses a large database of calibrated natural stereo-images with precisely co-registered laser-based distance measurements. First, we develop a procedure for precisely sampling stereo-image patches from the stereo-images, based on the distance measurements. The local depth variation in each stereo-image patch is quantified by disparity contrast. Next, we show that increased disparity contrast degrades performance in half-occlusion detection and disparity detection tasks, and changes the size and shape of the optimal spatial integration areas (“receptive fields”) for computing the task-relevant decision variables. Then, we show that a simple binocular image statistic predicts disparity contrast in natural scenes. Finally, we report results on the most likely patterns of disparity variation in natural scenes. Our findings motivate computational and psychophysical investigations of the mechanisms that underlie disparity estimation in local regions of natural scenes.

## INTRODUCTION

An ultimate goal of perception science and systems neuroscience is to understand how sensory-perceptual processing works in natural conditions. Unfortunately, natural stimuli are complex and difficult to characterize mathematically. In recent years, interest has increased in using natural stimuli for computational, psychophysical, and neurophysiological investigations (Adams et al., 2016; Burge & Geisler, 2011; 2012; 2014; 2015; Burge & Jaini, 2017; Burge, Fowlkes, & Banks, 2010; Burge, McCann, & Geisler, 2016; Cooper & Norcia, 2015; Felsen & Dan, 2005; Field, 1987; Geisler & Perry, 2009; Geisler & Ringach, 2009; Geisler, Najemnik, & Ing, 2009; Hibbard, 2008; Liu, Bovik, & Cormack, 2008; Maiello, Chessa, Solari, & Bex, 2014; Olshausen & Field, 1996; Potetz & Lee, 2003; Scharstein & Szeliski, 2003; Sebastian, Burge, & Geisler, 2015; Sprague, Cooper, Tosic, & Banks, 2015; van Hateren & van der Schaaf, 1998; Wilcox & Lakra, 2007; Yang & Purves, 2003). This burgeoning interest has been fueled by at least three factors. First, high-fidelity natural stimulus databases are becoming available for widespread scientific use. Second, powerful statistical, computational, and psychophysical methods are making natural stimuli increasingly tractable to work with. Third, and most importantly, the science requires it. Models of sensory and perceptual processing, from retina to behavior, that predict neurophysiological and behavioral performance with artificial stimuli often generalize poorly to natural stimuli. High-quality measurements of natural scenes and images are needed to ground experiments and models in the data that visual systems evolved to process.

The process by which visual systems estimate the three-dimensional structure of the environment is one of the most intensely studied questions in vision. The paradigmatic depth cue is binocular disparity. Stereopsis is the perception of depth based on binocular disparity, our most precise depth cue. In the vision community, stereopsis and the estimation of binocular disparity (i.e. solving the correspondence problem) have been investigated primarily with artificial images (but see (Burge & Geisler, 2014; Hibbard, 2008)). Researchers are developing psychophysical paradigms for using natural stimuli to investigate stereopsis, and computational analyses for uncovering the disparity processing mechanisms that optimize performance. Several natural stereo-image databases, some of which are accompanied by groundtruth distance measurements, have been released in recent years (Adams et al., 2016; Burge et al., 2016; Canessa et al., 2017; Scharstein & Szeliski, 2002). Research with natural stimuli is aided by methods for assigning accurate groundtruth labels to sampled stimuli. Sampling accuracy and precision must be at or above the precision of the human visual system. Otherwise, it is impossible to determine whether observed performance limits are due to inaccuracies in the sampling procedure or whether they are due to the properties of natural stimuli or the human visual system.

The primary aim of this manuscript is to determine the impact of local depth variation on half-occlusion detection and disparity detection performance, two tasks fundamentally related to stereopsis. These tasks are equivalent to i) determining whether a given point in one eye’s image has or lacks a corresponding point in the other eye’s image (i.e. half-occlusion detection) and ii) if it is binocularly visible, whether the other eye is foveating the same scene point as the first (i.e. disparity detection). Accurate performance in these tasks supports perception of depth order, da Vinci stereopsis, and fine stereo-depth discrimination (Blakemore, 1970; Cormack, Stevenson, & Schor, 1991; Kaye, 1978; Nakayama & Shimojo, 1990; Wilcox & Lakra, 2007). First, we develop a high-fidelity procedure for sampling stereo-image patches from natural stereo-images; we estimate that the procedure is as precise as the human visual system all but the most sensitive conditions (Blakemore, 1970; Cormack et al., 1991). (A Matlab implementation of the procedure is available at http://www.github.com/BurgeLab/StereoImageSampling.) Second, we show that local depth variation systematically degrades performance in both tasks, and changes the size and shape of the integration area that optimizes performance in both tasks. Finally, we examine how luminance and disparity vary in natural scenes and show how local depth variation can be simply estimated from stereo-images.

## RESULTS

To analyze the impact of natural depth variation on half-occlusion detection and binocular disparity detection in natural scenes, it is useful to sample a large collection of binocular image patches with groundtruth depth information. In natural stereo-images, groundtruth information about the 3D-coordinates of the imaged surfaces is typically unavailable. In computer graphics generated scenes, groundtruth information about the 3D scene is usually available. Unfortunately, it is unknown whether computed generated scenes accurately reflect all task-relevant aspects of natural images. Therefore, it is important to obtain natural stereo-image databases accompanied by the 3D-coordinates of each imaged surface. Provided the 3D natural scene data is of sufficiently high quality, groundtruth binocular disparities (and corresponding points) can be computed from the 3D data using trigonometric instead of image-based methods.

Recently, Burge et al. (2016) published a large database of calibrated stereo-images of natural scenes with precisely co-registered (+/-1 pixel) laser measurements of the groundtruth distances to the imaged objects in the scene. During acquisition of each eye’s view of the scene, the camera and the range scanner nodal points were positioned at identical locations. This feature of the data acquisition process is critical to ensure that each pixel in each eye’s photographic image has a matched pixel in associated range image, and vice versa. The current manuscript uses this dataset.

### Interpolating binocular corresponding points from groundtruth distance data

Accurate, precise determination of corresponding points is necessary for accurate, precise sampling of binocular image-patches. Corresponding image points are the left- and right-eye images of the same point in a scene. In natural stereo-images, corresponding image points are usually estimated via image-based methods (e.g. local cross-correlation). From the Burge et al (2016) dataset, it is possible to determine groundtruth corresponding points directly from the laser-measured distance data. It is also possible to determine, without resorting to image-based matching routines, whether a given point in one eye’s image has, or lacks, a corresponding point in the other eye’s image.

To obtain binocular image patches such that the center pixel of each eye’s patch coincides with a corresponding image points, a two-stage interpolation procedure is required. First, corresponding image point locations are interpolated using ray-tracing techniques. Second, the luminance and range images are interpolated to obtain stereo-image patches in which the center pixels of the left- and right-eye images coincide with corresponding image point locations. (Readers who are more interested in the paper’s scientific findings than in the methodological details of the stereo-image sampling and interpolation procedure can safely skip to the section “Quantifying local depth variation with disparity contrast”.)

Left- and right-eye image points are *corresponding image points* if they image the same surface point in a 3D scene. Sampling a pixel center from either the left- or the right-eye luminance image initializes the interpolation procedure. The eye from whose image the pixel center is first chosen is the *anchor eye*. Each pixel is located in a frontoparallel projection plane 3 meters from the cyclopean eye (i.e. midpoint of the interocular axis). Left-eye (LE) and right-eye (RE) lines of sight through the centers of these pixels define a set of intersection points in 3D space (Fig. 1A). These intersection points are the *sampled 3D scene points*. When a 3D surface in the scene passes through a sampled 3D scene-point, the left- and right-eye lines of sight to this point intersect the projection plane at pixel centers (Fig. 1A). However, most sampled 3D scene points do not have a 3D surface passing through them, and most 3D surface points do not coincide with sampled 3D scene points. Thus, corresponding image points do not generally coincide with pixel centers in the projection plane. The goal of our interpolation procedure is to interpolate 3D surface points and corresponding image points so that post-interpolation pixel centers coincide with corresponding image points

**Fig 1:**
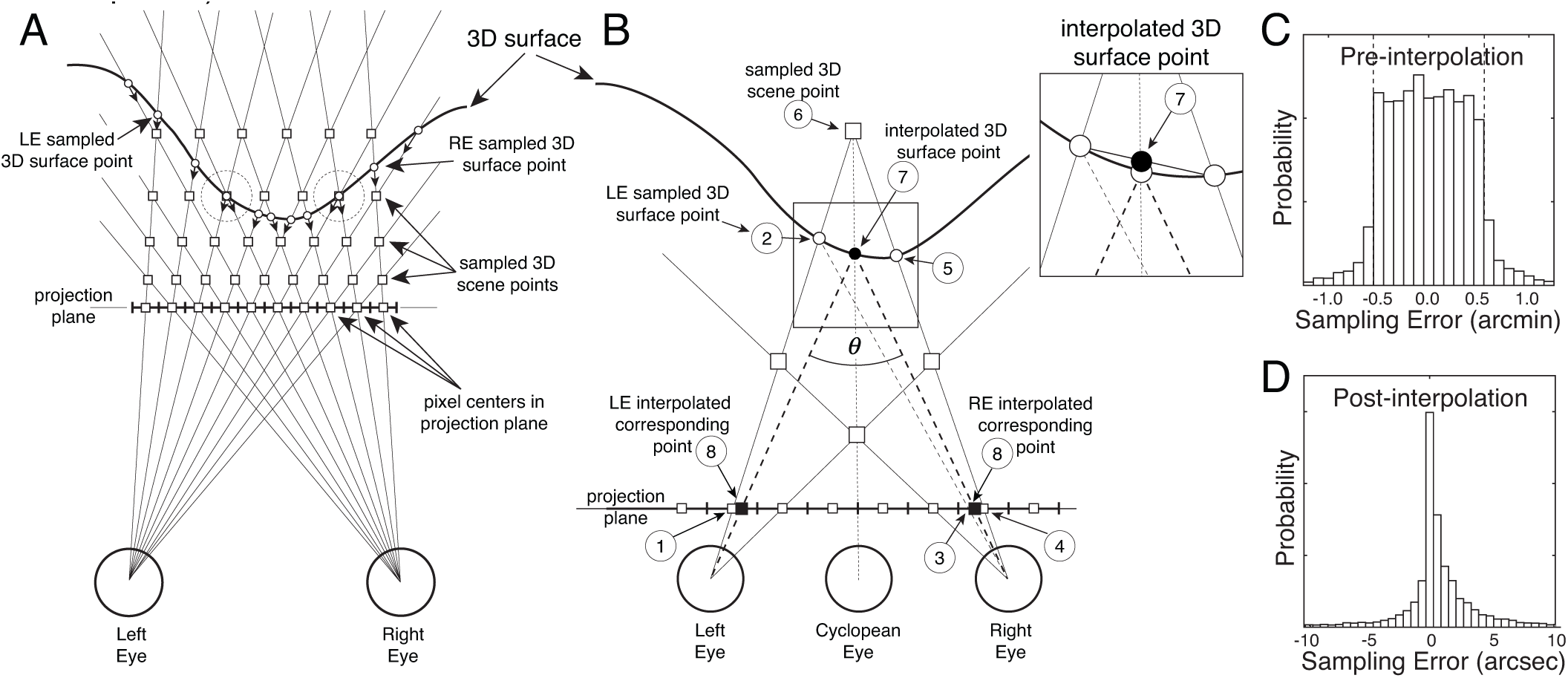
Stereo 3D sampling geometry, corresponding image-points, and interpolation procedure. **A** Top-view of 3D sampling geometry. Left-eye (LE) and right-eye (RE) luminance and range images are captured one human interocular distance apart (65 mm). Sampled 3D scene points (white squares) occur at the intersections of LE and RE lines of sight (thin lines) and usually do not lie on 3D surfaces. Samples in the projection plane (i.e. pixel centers) are a subset of these sampled 3D scene points. Sampled 3D surface points (white dots) occur at the intersections of LE or RE lines of sight with 3D surfaces (thick black curve) in the scene. Small arrows along lines of sight represent light reflected from sampled 3D surface points that determine the pixel values in the luminance and range images for each eye. Occasionally, sampled 3D surface points coincide with sampled 3D scene points (large dashed circles). Light rays from these points intersect the projection plane at pixel centers. **B** Procedure to obtain corresponding image point locations: (i) Sample a pixel location (1) in the anchor eye’s image (here, the left eye); (ii) Locate the corresponding sampled left eye 3D surface point (2); (iii) Find the right eye projection (3) from sampled 3D surface point by ray tracing. (iv) Select nearest pixel center (4) in right eye image; (v) Locate the corresponding sampled right eye 3D surface point (5). (vi) Find sampled 3D scene point (6) nearest the left- and right-eye sampled 3D surface points. This sampled 3D scene point is the intersection point of the left- and right-eye lines of sight through the sampled 3D surface points. (vii) Find interpolated 3D surface point (7) by linear interpolation (i.e. the location of the intersection of cyclopean line of side with chord joining sampled 3D surface points; see inset); (viii) Dashed light rays from this interpolated 3D surface point define corresponding point locations (8) in the projection plane. The vergence demand *θ* of the interpolated scene point is the angle between the left- and right-eye lines of sight required to fixate the point. **C** Sampling error before interpolation in arcmin. Dashed vertical lines indicate the expected sampling error assuming surface point locations are uniformly distributed between sampled 3D scene points. **D** Estimated sampling error after interpolation in arcsec.

Fig. 1B illustrates how interpolated 3D surface and corresponding point locations are obtained. Consider a pair of LE and RE pixel centers that correspond to a sampled 3D scene point. Sampled 3D scene points (Fig. 1B, open squares in scene) do not generally coincide with sampled 3D surface points (Fig. 1B, open circles). Thus, the luminance information in these pixels (Fig. 1B, open squares in projection plane) does not generally correspond to a single point on a 3D surface. The interpolated surface point (Fig. 1B, black circle) occurs at the intersection between the cyclopean line of sight and a line segment connecting sampled 3D surface points. This interpolated 3D surface point, unlike the sampled 3D scene point, lies on (or extremely near to) a 3D surface. The LE and RE lines of sight to the interpolated 3D surface point intersect the projection plane at *corresponding image points* (Fig. 1B, black squares).

This interpolation procedure is necessary to ensure to that binocular sampling errors are below human disparity detection thresholds. Under optimal conditions, human disparity detection thresholds are approximately 5 arcsec (Blakemore, 1970; Cormack et al., 1991). Failing to interpolate would result in ±35arcsec binocular sampling errors (i.e. erroneous fixation disparities), which are large relative to disparity detection thresholds. Assuming surfaces are uniformly distributed between sampled 3D scene points, the difference between the vergence demand of the interpolated 3D surface point and the nearest 3D sampled scene point should be uniformly distributed. (Vergence demand *θ* is the angle between the LE and RE lines of sight required to fixate the scene point.) Figure 1C confirms this prediction; the vergence demand differences indeed tend to lie between ±35arcsec, indicating that the assumptions of the interpolation procedure are accurate.

Unfortunately, interpolated image points returned by this procedure are not guaranteed to be true corresponding points. If a sampled surface point is half-occluded, for example, corresponding image points do not exist, so the procedure returns points that are invalid. Invalid points can also occur if the distance data is corrupted by noise. We screen for bad points by repeating the interpolation procedure twice, with a different eye serving as anchor eye on each repeat. When the interpolated 3D surface points and their associated vergence demands match on both repeats, it indicates that the interpolated corresponding points are valid. Figure 1D shows that after interpolation, approximately 80% of interpolated 3D surface points had vergence demand differences of less than ±5 arcsec across repeats. For subsequent analyses of binocularly visible scene points, interpolated points with vergence demand differences larger than ±5 arcsec are discarded, ensuring that residual sampling errors are smaller than human stereo-detection thresholds for all but the very most sensitive conditions (Blakemore, 1970; Cormack et al., 1991). Visual inspection of hundreds of interpolated points corroborates the numerical results (see below).

To understand why differences in vergence demand can help identify half-occluded points, consider the scenario depicted in Fig. 2A. When the left eye is the anchor eye, the left-eye image point is associated with a far surface point having vergence demand *θ* _*L*_, and the right-eye image point returned by the interpolation is invalid because no true corresponding point exists. When the right eye is the anchor eye, the same right-eye image point is associated with a near surface point having vergence demand *θ* _*R*_. In this scenario, the vergence demand difference Δ*θ* = *θ* _*R*_ - *θ* _*L*_ clearly does not equal zero. Also note that when the right eye is the anchor eye, the left-eye image point (middle black square) returned by the procedure does not match the original left-eye image point. For cases in which the surface point is binocularly visible, both repeats of the interpolation procedure yield the same vergence demands, surface points, and interpolated image points (Fig. 2B). The vergence demand of a surface point is computed in the epipolar plane defined by the surface point and the left- and right-eye nodal points (Fig. 2C).

**Figure 2.**
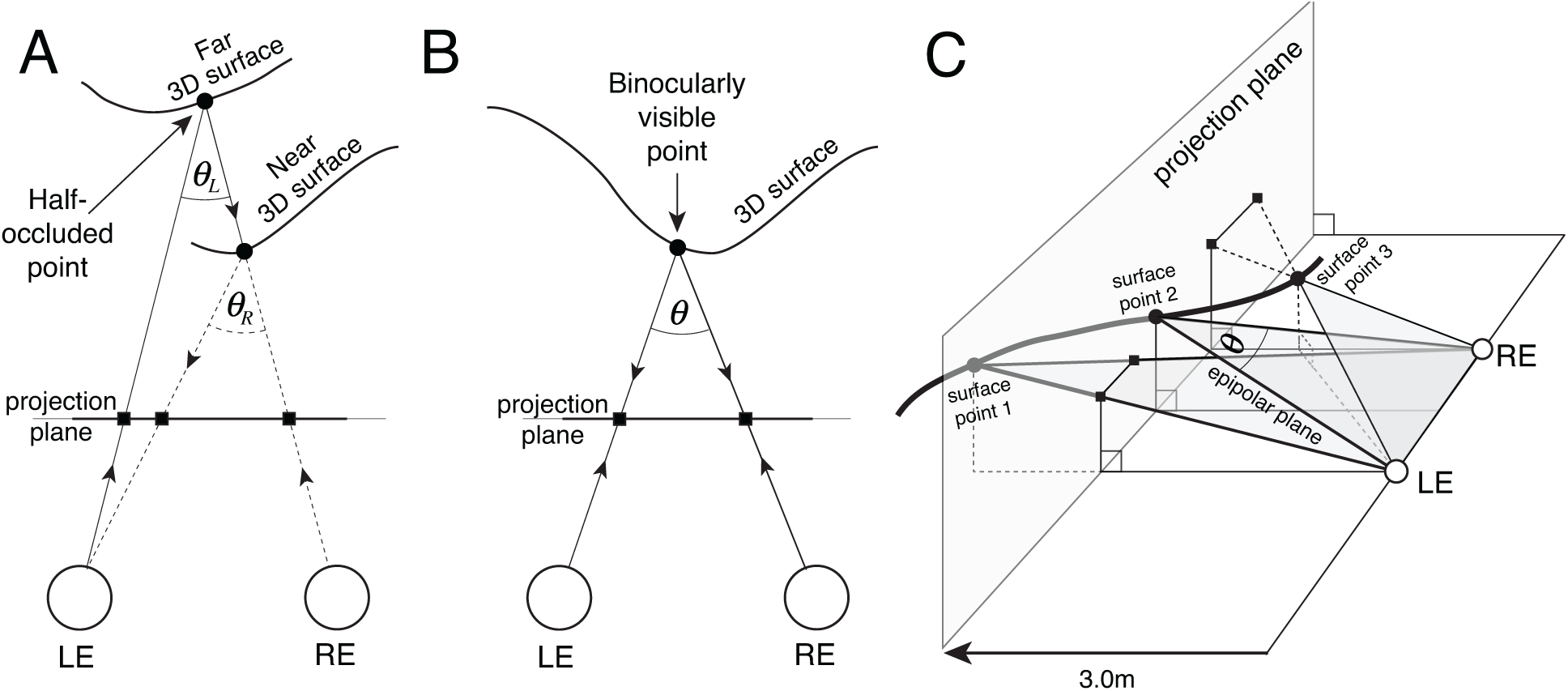
Half-occluded scene points, binocularly visible scene points, and vergence demand. **A** Half-occluded 3D surface point. The scene point on the far surface (black circle) is visible to the left eye and occluded from the right eye. Arrows indicate the ray tracing performed by the interpolation routine (see Fig. 1). Squares represent interpolated image points returned by the interpolation procedure. When 3D surface points are half-occluded, the interpolation procedure returns invalid points. **B** Binocularly visible surface point (black circle) and corresponding image points (black squares) in the projection plane. When the scene point is binocularly visible, the vergence demand *θ* of the surface point is the same, regardless of the anchor eye. The vergence demand is identical whether the left or the right eye is used as the anchor eye. **C** Vergence demand is computed within the epipolar plane defined by a 3D surface point and the left- and right-eye nodal points.

Results of the sampling and interpolation procedure are depicted in each of two natural scenes (Fig. 3A,B). Left- and right-eye photographic images (upper row) and range images (lower row) are shown. 500 randomly sampled corresponding image points, associated with 500 scene points, are overlaid onto each stereo-image; 250 were sampled with the left eye as the anchor eye and 250 were sampled with right eye as the anchor eye. Divergently-fuse the left two images or cross-fuse the right two images to see the scene and the corresponding points in stereo-3D. True corresponding image points (yellow) lie on the images surfaces in the three-dimensional scene. Note that invalid interpolated points (red) are also shown. To protect against eye-specific biases in the subsequent analyses, surface points are sampled symmetrically about the sagittal plane of the head.

**Figure 3.**
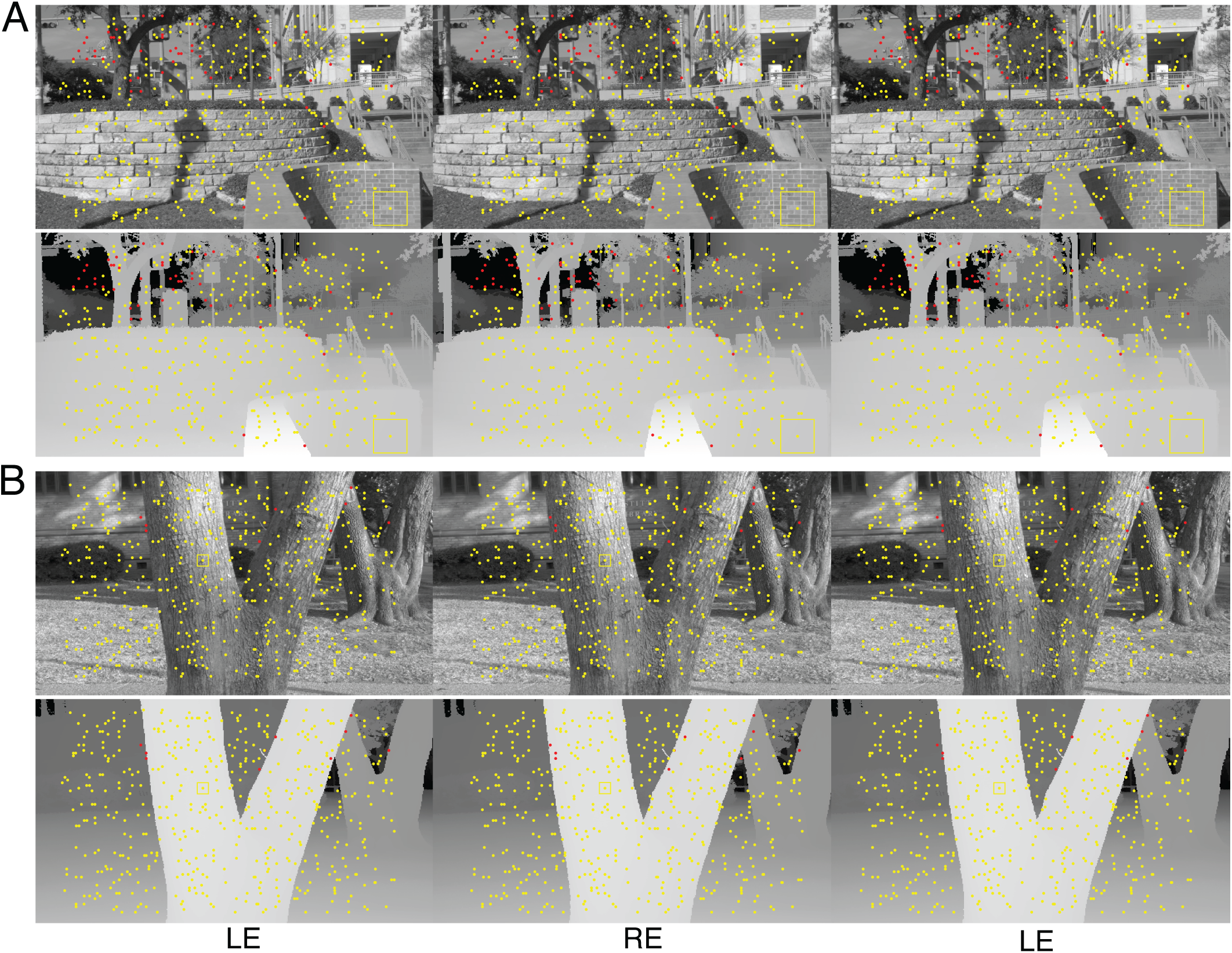
Corresponding points overlaid on stereo-images (upper row) and co-registered groundtruth distance data (lower row) for two different scenes (**A** and **B**). Wall-fuse the left two images or cross-fuse the right two images to see the imaged scene in stereo-3D. True corresponding points (yellow dots) lie on imaged 3D surfaces. Candidate corresponding points that are half-occluded or are otherwise invalid (red dots) are also shown. For reference, the yellow boxes in A and B indicate 3° and 1° areas, respectively.

After corresponding points are determined, left- and right-eye luminance and range stereo-image patches centered on the corresponding points are cropped. Luminance and range values are interpolated on a uniform grid of pixel-centers centered on at the corresponding points. Maps of the groundtruth disparity, relative to the center pixel, can then be computed directly from the range images.

### Quantifying local depth variation with disparity contrast

The patterns of binocular disparities encountered by a behaving organism depend on the properties of objects in the environment and how the organism interacts with those objects. When an organism fixates a point on an object in a 3D scene, its image is formed on the left and right-eye foveas. These images are the inputs to the organism’s foveal disparity processing mechanisms. To first approximation, if the fixated point lies on a planar frontoparallel surface, then disparities of nearby points will be zero. However, when the fixated point lies on curved, bumpy, and/or slanted surface, the disparities of nearby points will vary more significantly. When a depth edge is near the fixated point, dramatic changes in disparity can occur in the neighborhood of the fovea.

To quantify local depth variation, we compute the disparity contrast associated with each stereo-pair that is centered on a binocularly visible scene point. Disparity contrast is the root-mean-squared (RMS) disparity relative to the center pixel in a local neighborhood

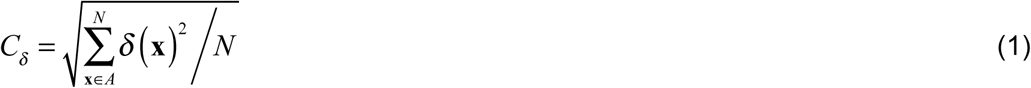

where *δ* is the groundtruth relative disparity, *A* is the local spatial integration area, **x** is the spatial location, and *N* is the total number of pixels in the local area. Example stereo-images with different amounts of disparity contrast are shown in Fig. 4. The upper row shows the luminance stereo image. The lower row shows the groundtruth disparity map, computed directly from the laser-measured distance data at each stereo-image pixel. Thus, each stereo-image patch corresponds to the left- and right-eye retinal image that would be formed if an observer fixated the surface point in the scene. The distribution of disparity contrast in natural scenes is shown in Fig. S1.

**Figure 4.**
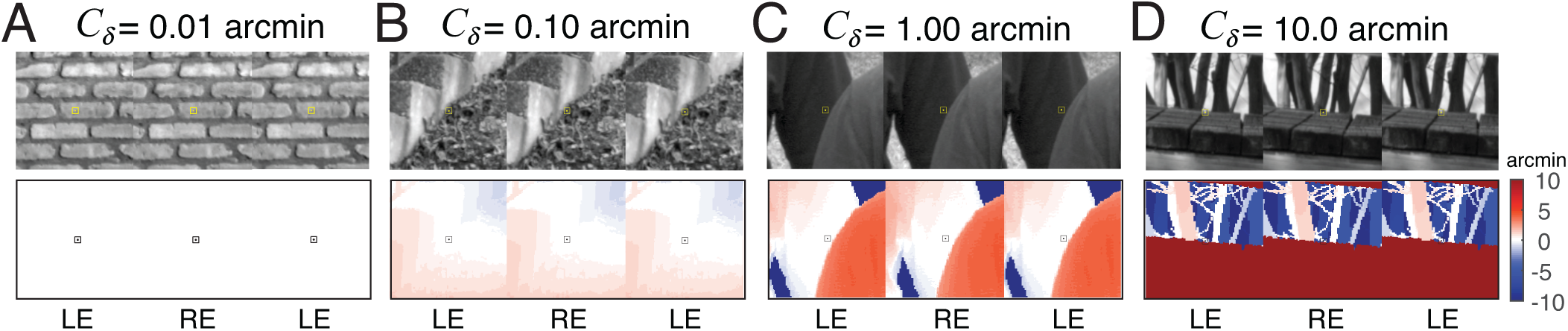
Natural stereo-image patches and corresponding groundtruth disparity maps, sampled from natural scenes. Free-fuse to see in stereo-3D. **A-D** Local disparity contrast *C*_*δ*_ (e.g. local depth variation) increases in the subplots from left to right. Groundtruth disparity at each pixel (bottom row) are computed directly from groundtruth distance data. Disparity contrast is computed under a window that defines the spatial integration area (see Methods). The colorbar indicates the disparity in arcmin relative to the disparity (i.e. vergence demand) at the center pixel.

### Half-occlusion detection in natural stereo-images

Half-occlusion detection is the task of detecting if a particular scene point visible to one eye is occluded to the other eye. This task is equivalent to determining whether a given point in one eye’s image lacks or has a corresponding point in the other eye. Half-occlusion detection is important because disparity is defined only when a given point is binocularly visible, and because half-occluded points can mediate da Vinci stereopsis (Harris & Wilcox, 2009; Kaye, 1978; Nakayama & Shimojo, 1990).

What image cues provide information about whether scene points are half-occluded or binocularly visible, and how does local depth variation impact the information? First, consider half-occluded scene points (Fig. 5A). If the eyes are verged on (i.e. pointed towards) a half-occluded point (see Fig. 2A), the scene point at the center of one eye’s image is different than the scene point at the center of the other eye’s image, the left- and right-eye images will be centered on different points in the scene, and the left- and right-eye images should be very different (Fig. 5B). Now, consider binocularly visible scene points. If the eyes are verged (i.e. fixated) on a binocularly visible scene point, the left- and right-eye images should be very similar. However, if local depth variation near a binocularly visible scene point is high, left- and right-eye images centered on that point should be less similar.

**Figure 5.**
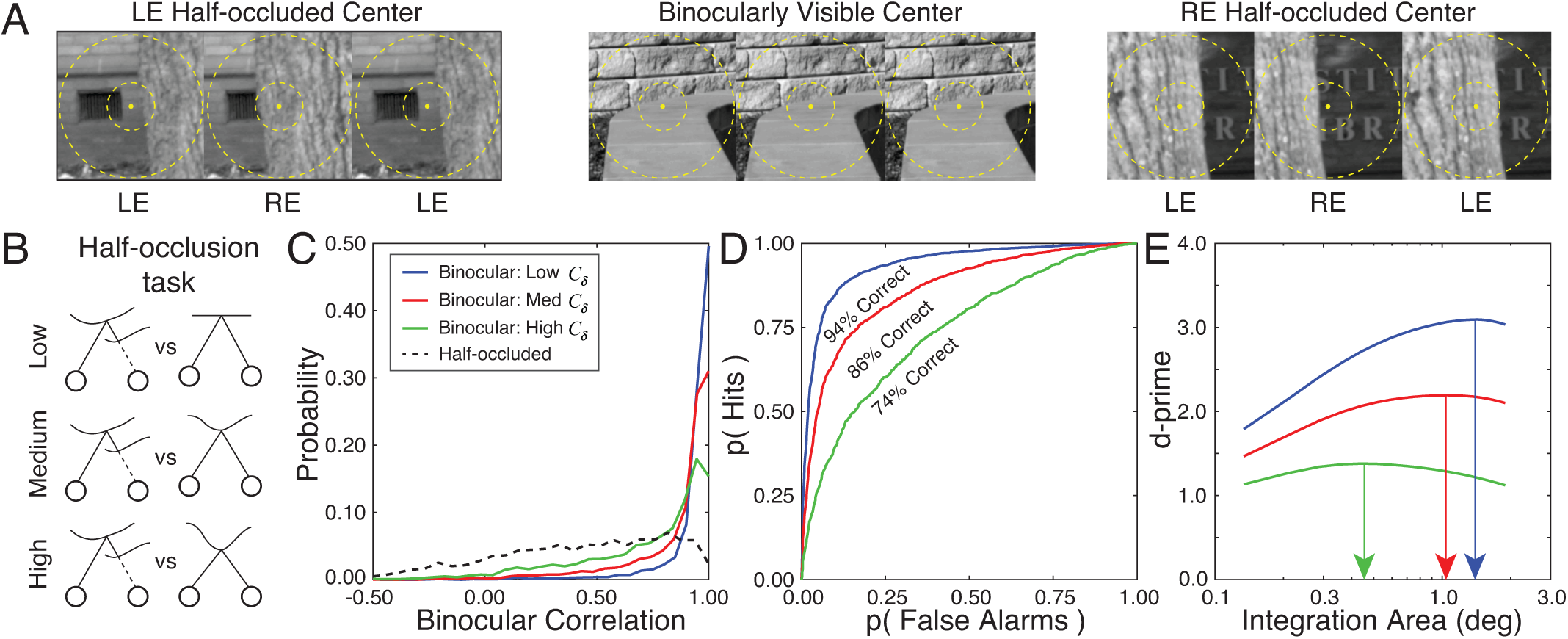
Effect of disparity contrast on half-occlusion performance. **A** Three example stereo-image patches centered on scene points that are half-occluded to the left eye, binocularly visible, and half-occluded to the right eye. Spatial integration areas of different sizes (1° and 3°) are shown as dashed circles. **B** The half-occlusion detection task is to distinguish half-occluded vs. binocularly visible points with 0.0 arcmin of disparity. Performance is compared for scene points with low, medium, and high disparity contrasts. **C** Conditional probability distributions of the decision variable (i.e. the binocular correlation of the left- and right-eye image patches). The dashed black curve represents the distribution of the decision variable for half-occluded points. Solid curves show the decision variable distributions for patches with binocularly visible centers having low (blue; 0.05-1.00 arcmin), medium (red; 0.2-4.0 arcmin), and high (green; 0.75-15.0 arcmin) disparity contrasts. Binocular image correlation and disparity contrast are computed with spatial integration areas of 1.0° (i.e. 0.5° at half-height). **D** Receiver operating characteristic (ROC) curves for the half-occlusion task. Higher disparity contrasts decrease half-occlusion detection performance. **E** Half-occlusion detection sensitivity (d-prime) as a function of spatial integration area for different disparity contrasts. Arrows mark the spatial integration area at half-height for which half-occlusion detection performance is optimized.

To examine the impact of local disparity variation on half-occlusion detection in natural scenes, we performed the following analysis. First, we sampled 10,000 of stereo-image patches from the natural scene database using the methods discussed above. 86.5% of the sampled stereo-image pairs were centered on binocularly visible scene points; 13.5% were centered on half-occluded scene points. We determined which patches had half-occluded centers directly from the range measurements by determining which patches had centers where the horizontal disparity gradient (*DG* = Δ' Δ*X*) with any other point was 2.0 or higher. Second, to quantify local depth variation, we computed the disparity contrast of all patches with binocularly visible centers. For all analyses, disparity contrast was computed over a local integration area of 1.0° (0.5° at half-height; see Eq. 1); results are robust to this choice. Third, the similarity of the left- and right-eye image patches was quantified with the correlation coefficient

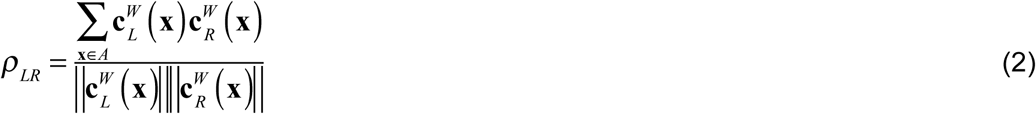

Where 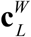 and 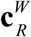 are windowed left- and right-eye Weber contrast images (see Methods) and where 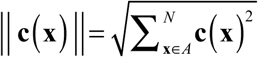 is the L2 norm of the contrast image in a local integration area *A*. Importantly, the window determines the size of the spatial integration area within which binocular correlation is computed. Fourth, under the assumption that the correlation coefficient is the decision variable, we used standard methods from signal detection theory to determine how well half-occlusions can be detected in natural images. Specifically, we determined the conditional probability of the decision variable given i) that the center pixel was binocularly visible for each disparity contrast *p*(*p*_*LR*_ | *bino,C*_*δ*_) and ii) that the center pixel was half-occluded *p*(*p*_*LR*_ | *mono*) (Fig. 5C), swept out an ROC curve (Fig. 5D), computed the area underneath it to determine percent correct, and then converted to d-prime. Finally, we repeated the steps as a function of the spatial integration area. Half-occlusion detection performance (d-prime) changes significantly as a function of the spatial integration area for each of several disparity contrasts (Fig. 5E).

The results in Figure 5D,E show that local depth variation reduces how well binocularly visible points can be discriminated from half-occluded points. Figure 6A summarizes half-occlusion detection performance with the best integration area, for more finely spaced bins of disparity contrast (also see Fig. S2A). Figure 6B summarizes how increasing disparity contrast decreases the size of the spatial integration area that optimizes performance. Compared to when the best-fixed integration area is used across all stimuli, d-prime is 8% higher when the optimal spatial integration area is used for each stimulus (see Methods). Thus, in half-occlusion detection, the visual system would benefit from half-occlusion detection mechanisms, should adapt its spatial integration area to the local depth variation in the scene.

**Figure 6.**
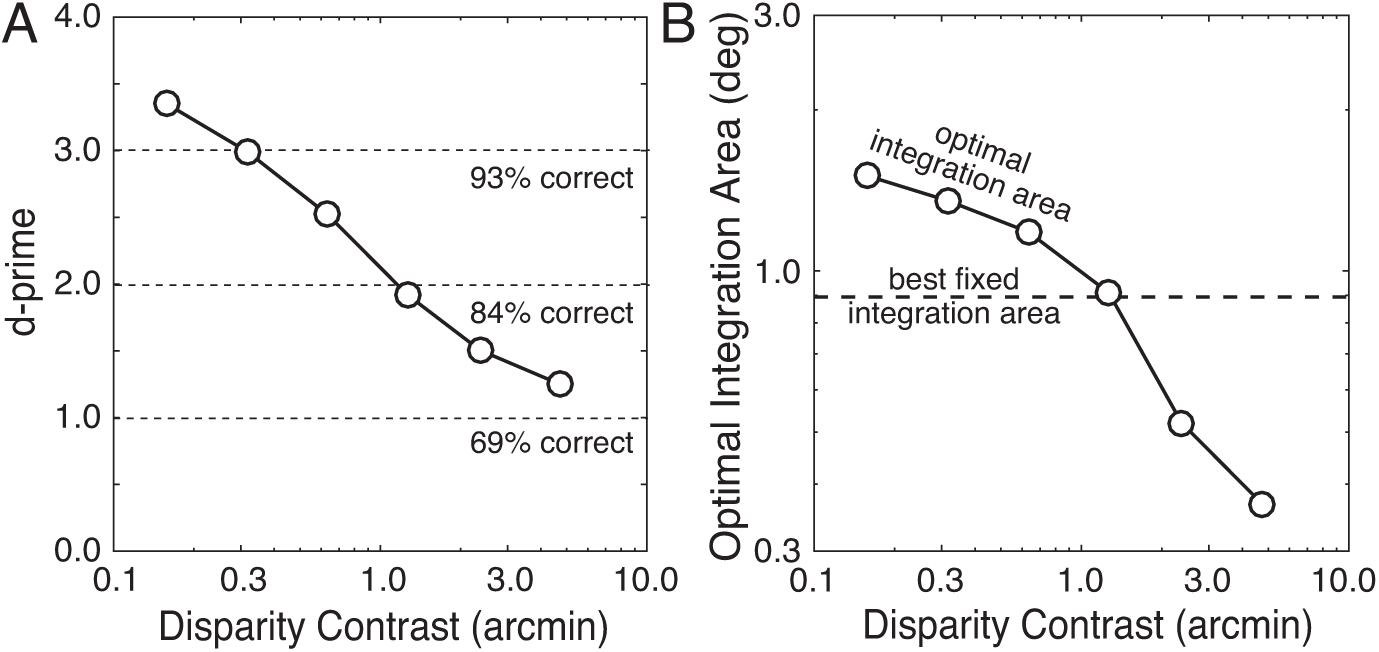
Effect of local disparity variation on optimal processing size for half-occlusion detection. **A** Sensitivity as a function of disparity contrast, assuming the optimal size of the integration area. Sensitivity decreases monotonically with disparity contrast. For each disparity contrast, sensitivities were measured with the optimal integration area. **B** Optimal window size as a function of disparity contrast. The optimal window size decreases approximately linearly as disparity contrast increases on a log-log scale. Results are highly robust to changes in the bin width.

### Disparity detection in natural stereo-images

Binocular disparity is our most precise depth cue. Binocular disparity detection is the task of detecting whether a particular binocularly visible point is being perfectly foveated (i.e. fixated) or not. When a target point is fixated accurately, the point is imaged on the foveas of the two eyes. When a target point is not fixated accurately, the point’s image will not fall on the foveas and non-zero disparities occur (Fig. 7A). Just as local depth variation impacts the ability to detect whether a point in one eye’s image is half-occluded or binocularly visible, local depth variation should impact the detection of non-zero binocular disparities in natural scenes (Fig. 7B).

**Figure 7.**
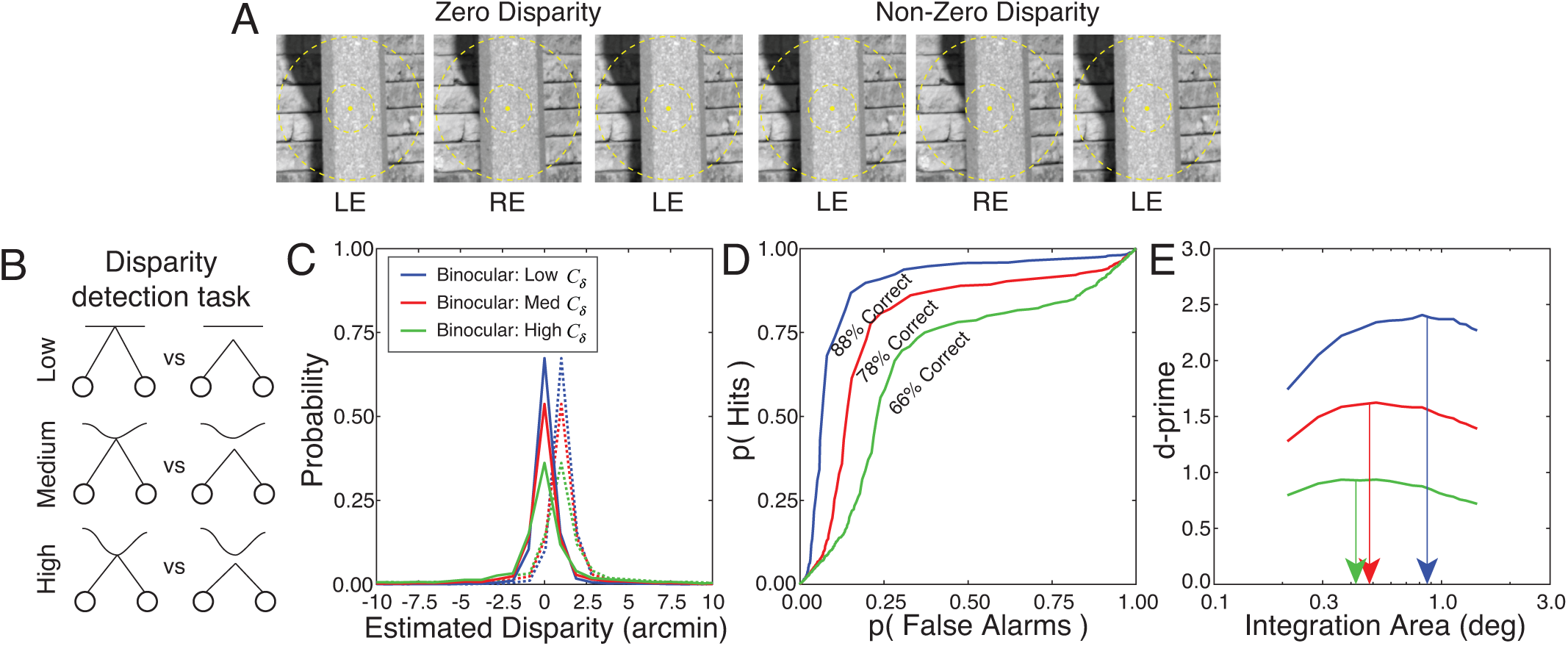
Effect of disparity contrast on disparity detection performance. **A** Stereo-image patches centered on binocularly visible scene points with 0.0 and 1.0 arcmin of fixation disparity. The eyes are fixated 1 arcmin in front of the target in the right image. **B** The disparity detection task simulated here is to distinguish scene points with 0.0 arcmin vs. 1.0 arcmin of fixation disparity. Performance is compared for scene points with low, medium, and high disparity contrasts. **C** Conditional probability distributions of the decision variable. The decision-variable is the disparity that maximizes the local cross-correlation function (Eq. 3). Results are presented for a spatial integration area of size 1.0°. The solid and dashed curves show the decision variable for scene points fixated with 0.0 and 1.0 arcmin of disparity, respectively, for patches having low (blue), medium (red), and high (green) disparity contrasts. **D** ROC curves for disparity detection. **E** Disparity detection sensitivity (i.e. d-prime) as a function of spatial integration area for different disparity contrasts.

Local windowed cross-correlation is the standard model of disparity estimation (Banks, Gepshtein, & Landy, 2004; Cormack et al., 1991; Tyler & Julesz, 1978). Under this model, the estimated disparity is the disparity that maximizes the inter-ocular correlation between a reference patch in one eye’s image and a test patch in the other eye’s image.

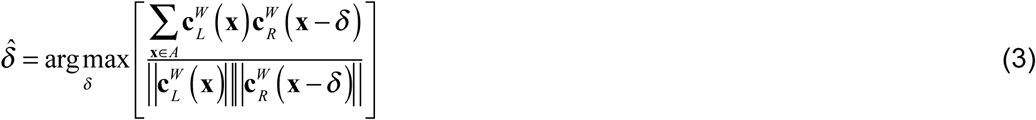

where 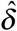 is the disparity estimate and *δ* is the disparity between a patch in the anchor eye and a patch in the other eye. Equation 3 is written assuming that the left eye anchor eye.

To examine the impact of local depth variation on disparity detection thresholds, we performed an analysis nearly identical to the half-occlusion detection analysis presented above. First, we randomly sampled 10,000 stereo-image patches having zero absolute disparity at the center pixel. Second, we estimated disparity from the stereo-image patches using local windowed cross-correlation (Eq. 3). These disparity estimate is the decision variable for the disparity detection task. The conditional probability of the disparity estimates for each disparity contrast 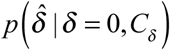 is symmetric and centered at zero (Fig. 7C). Third, assuming that the distribution of estimates for small non-zero disparities 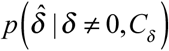 is a shifted version of the distribution for zero disparities, we swept out an ROC curve (Fig. 7D), computed the area underneath it to determine percent correct, and then converted to d-prime. Finally, for each disparity contrast, we computed d-prime for detecting a target with 1.0 arcmin of disparity as a function of different local integration areas (Fig. 7E).

Results for the disparity detection task are similar to results for the half-occlusion task. Local depth variation reduces disparity detection sensitivity (Fig. 7D,E; Fig. 8A; Fig. S2B), and decreases the size of the spatial integration area that optimizes performance (Fig. 7E; Fig. 8B; Fig. S2B). When the integration area is too large, the depth variation within the integration area prevents reliable estimates. When the integration area is too small, the luminance variation within the integration area is insufficient to obtain a reliable estimate. Thus, the visual system should adapt its spatial integration area to the local depth variation in the scene. Compared to when the best-fixed integration area is used across all stimuli, d-prime is 12% higher when the optimal spatial integration area is used (see Methods). Unlike half-occlusion detection, however, the optimal integration area for disparity detection shrinks and then plateaus (Fig. 8B), and does not decrease below 0.4° (0.2° at half-height).

**Figure 8.**
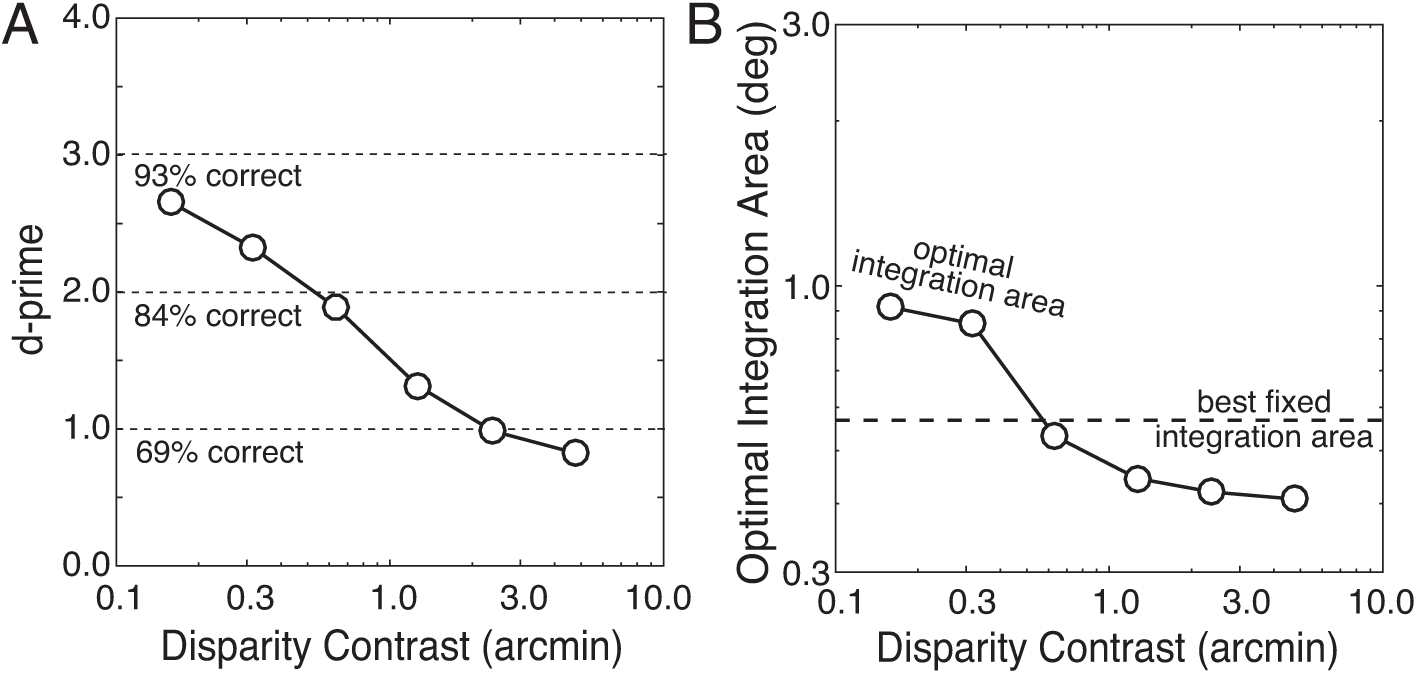
Effect of local disparity variation on size of optimal integration area for disparity detection. **A** Sensitivity as a function of disparity contrast. Sensitivity drops monotonically with disparity contrast. For each disparity contrast, sensitivities were measured with the optimal integration area. **B** Optimal integration area as a function of disparity contrast. The optimal integration area decreases as disparity contrast increases and then plateaus at a minimum value (0.4° window; 0.2° at half-height).

Interestingly, these results are closely related to results in the psychophysics literature on human stereopsis. Local depth variation hurts human performance in depth discrimination, disparity detection, and stereo-resolution tasks (Banks et al., 2004; Ernst & Banks, 2002; Harris, McKee, & Smallman, 1997; Kane, Guan, & Banks, 2014). (Spatial stereo-resolution tasks measure the finest detectable spatial modulation of binocular disparity.) Two separate groups have argued that human spatial stereo-resolution is limited by the smallest disparity selective receptive fields available to the human visual system (Banks et al., 2004; Harris et al., 1997). Harris et al. estimated that the smallest disparity-selective receptive fields available to the human visual system are 0.07-0.13° in diameter (Harris et al., Optimal Integration Area (deg) 1997). Banks et al. estimated that the smallest receptive fields are 0.13° in diameter (Banks et al., 2004).

Our estimate of the smallest useful disparity receptive field in natural scenes (0.4° integration area, 0.2° width at half-height) is within a factor of two to the psychophysical estimates of the smallest receptive field available to the human visual system (~0.1°). Thus, just as the sampling rate of the foveal cone photoreceptors is determined by the cut-off spatial frequency of human optical point spread function, the smallest disparity selective fields available to the human visual system may be determined by the smallest receptive fields that are useful for estimating disparity in natural binocular images. The logic is that there is little point in having receptive fields that select for information that is not useful or available.

### Effect of depth variation on optimal shape of integration region

The previous sections demonstrate that the optimal spatial integration area for half-occlusion and disparity detection decreases in size with increases in disparity contrast. Does disparity contrast also impact the shape of the optimal spatial integration areas? To check, we performed the following steps, starting with the half-occlusion task. First, for a given disparity contrast, we found the optimally sized integration area (see Figs. 6B, 8B). Second, we varied the aspect ratio of the integration area while holding the size of the integration area fixed (Fig. 9A), computed the task-relevant decision variable (Eq. 2) for each aspect ratio, and determined d-prime using the procedures described above. Third, we repeated the previous steps across different disparity contrasts and plotted sensitivity. Performance is optimized by vertically elongated integration areas at high disparity contrasts, and by (slightly) horizontally elongated areas at low disparity contrasts (Fig. 9B,C). We repeated these analysis for the disparity detection task and found that the same patterns hold (Fig. 9D-F). The secondary effect of optimizing aspect-ratio is modest (~0.1 in d-prime units) compared to the primary effect of optimizing size. However, given that evolution tends to push organisms towards the optimal solutions in critical tasks, one might expect biological systems to have developed mechanisms that adapt both the size and the shape of their receptive fields to the local depth structure of stimuli. Investigating whether visual systems have developed such mechanisms will be an interesting topic for future research.

**Figure 9.**
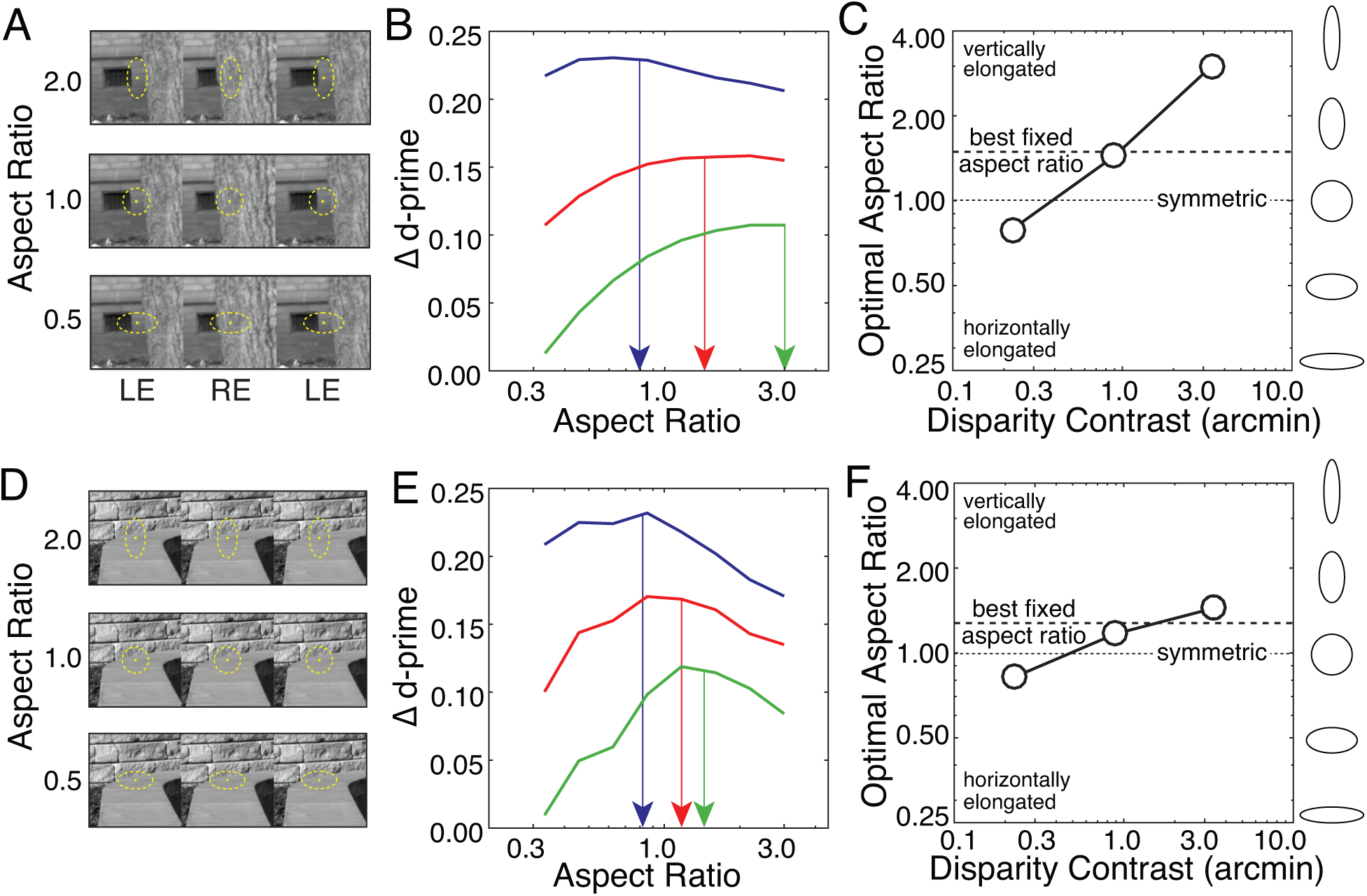
Effect of local depth variation on the shape of the spatial integration area that optimizes performance. **A** Integration areas with the same size but different aspect ratios within which to compute the decision variable for the half-occlusion task (i.e. binocular image correlation). **B** Change in half-occlusion detection sensitivity (i.e. d-prime) as a function of aspect ratio for different disparity contrasts. Arrows indicate the aspect ratio that maximizes half-occlusion detection performance. The maxima were determined using a polynomial fit (not shown) to the raw data. Aspect ratios less than 1.0 are horizontally elongated. Aspect ratios larger than 1.0 are vertically elongated. Colors indicate low (blue; 0.05-1.00arcmin), medium (red; 0.2-4.0arcmin), and high (green; 0.75-15.0arcmin) disparity contrasts **C** Optimal aspect ratio as a function of disparity contrast. The optimal window for half-occlusion detection is more vertically elongated for higher disparity contrasts. The best-fixed aspect ratio across all disparity contrasts is also shown. **D** Same as A, but for the disparity detection task. **E-F** Same as B-C, but for the disparity detection task.

Why does the shape of the optimal integration area change with disparity contrast? From visual inspection of numerous individual examples we speculate that, at high disparity contrasts, vertical elongation improves performance because large disparity contrasts are most often caused by vertically oriented depth edges (e.g. Fig. 9A). For such cases, vertically oriented integration areas increase the number of pooled spatial locations over which the disparity is more nearly constant (Kanade & Okutomi, 1994). We are less clear about why, at low disparity contrasts, integration areas with slight horizontal elongation improves performance. We speculate that this is because low disparity contrasts are often associated with the ground plane (e.g. Fig. 9D), and horizontally oriented integration areas maximize the number of pooled spatial locations with the same disparity.

### Estimating disparity contrast

Local depth variation in the region around fixation makes disparity-related tasks more difficult (Fig 5-8). A visual system with access to estimates of local disparity contrast can, in principle, improve half-occlusion and disparity detection performance by adapting the size and shape and shape of its receptive fields to local disparity contrast. How might the visual system estimate disparity contrast from information in the left- and right-eye images? One approach is to estimate disparity at each spatial location (pixel) in a local area with generic receptive fields, compute the contrast (i.e. local root-mean-squared disparity) of those estimates, and then re-estimate the disparities with optimized receptive fields. A second more direct approach is to compute a simple image statistic that predicts disparity contrast at each spatial location, and then estimate the disparities with optimized receptive fields.

Interestingly, the contrast *C*_*B*_ of the binocular difference image is a good predictor of disparity contrast *C*_*δ*_. The binocular difference image is the pixel-wise difference between the left- and right-eye Weber contrast images 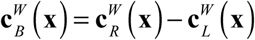. Figure 10A shows a stereo-image patch with low disparity variation and low binocular difference image contrast. Figure 10B shows a stereo-image patch with high disparity contrast and high binocular difference image contrast. Figure 10C shows that difference image contrast predicts disparity contrast across thousands of patches (n=10,000).

**Figure 10.**
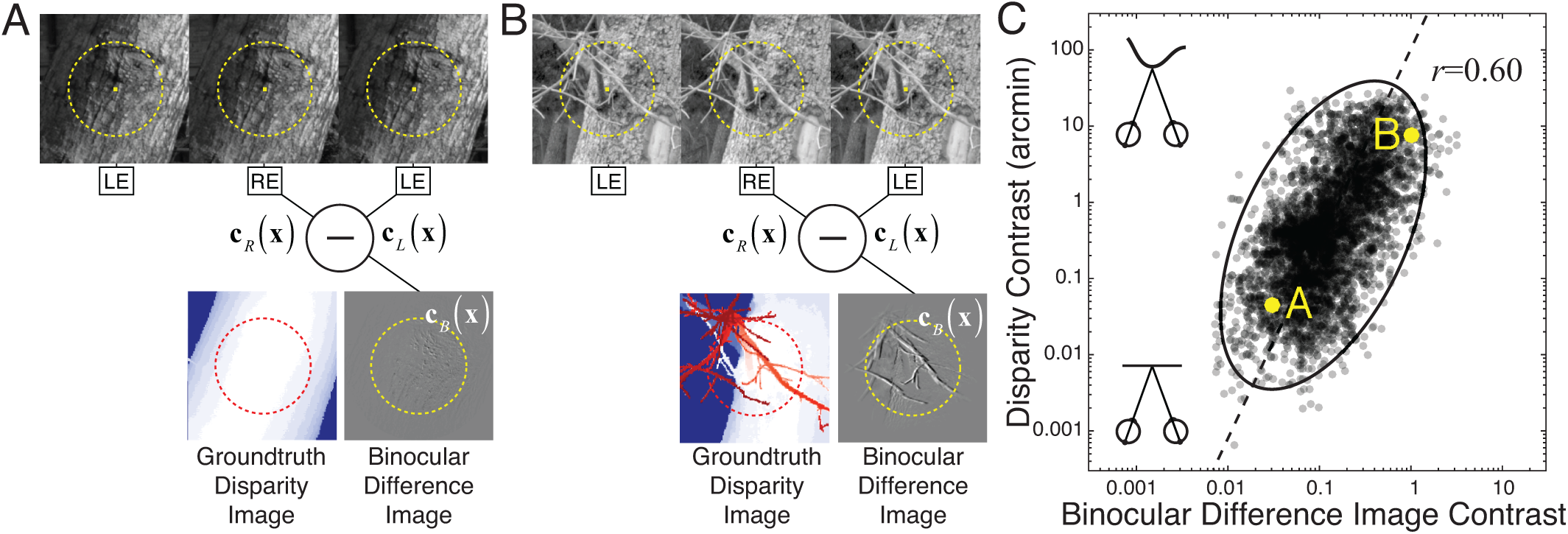
Joint statistics of disparity contrast and binocular difference image contrast in natural scenes. **A** Stereo-image with low groundtruth disparity contrast and low binocular difference image contrast. The upper row shows the stereo-image; the circle indicates the 1° spatial integration area from which the statistics were computed. The lower row shows the groundtruth disparities and the binocular difference image. **B** Stereo-image with high groundtruth disparity contrast has high binocular difference image contrast. **C** Disparity contrast and binocular difference image contrast in natural scenes are jointly distributed as a log-Gaussian and are significantly correlated. Points labeled in yellow indicate the disparity contrast and binocular difference image contrast of the stereo-images A and B. Statistics were computed for a spatial integration area of 1.0° (0.5° width at half-height). Similar results hold for other spatial integration areas.

Binocular difference image contrast and disparity contrast are jointly log-Gaussian distributed and are strongly correlated (*r* = 0.60; Fig. 10C). The correlation is nearly independent of viewing distance, although the most likely disparity contrasts decrease as distance increases. The relationship is nicely fit by a line in the log domain and a power law 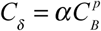 in the linear domain where *p* is the power and *α* is a proportionality constant (i.e. Weber Fraction); the best-fit power is *p* ≅ 2.0. Thus, the visual system could obtain a relatively precise estimate of local disparity variation (and local depth variation) directly from the local contrast of the binocular difference image, and use this estimate to select the integration area that is best-suited for a given level of disparity contrast (Chen & Qian, 2004). We believe these findings motivate a series of investigations on the mechanisms for estimating local disparity variation, and assessing its impact on disparity detection performance in natural scenes.

### Natural statistics of disparity variation

Local disparity variation negatively impacts half-occlusion and disparity detection performance. What is the most likely spatial pattern of disparity variation in natural scenes? Assuming that fixations occur only on 3D surface points, the disparity at the foveas is always zero. At non-foveal retinal locations, disparities vary with the depth structure (and distance) of the fixated stimulus. The standard deviation of disparity across multiple stimuli, at each retinal location, quantifies the most likely pattern of disparity variation.

Figure 11 shows how the pattern of disparity variation changes with retinal eccentricity near the fovea (±0.5°). Disparities are zero at the foveas, by definition, and become more variable at retinal positions farther from fixation. Interestingly, the region of least disparity variation is slightly vertically elongated (Fig. 11A). The impact of disparity contrast on this spatial pattern is shown in Fig. 11B. Spatial patterns of near-foveal disparities for stimuli with different amounts of local depth variation (i.e. disparity contrast) can be compared in Fig. 11B. Stimuli with low disparity contrast tend to have horizontally oriented regions of least disparity variation around the fovea. Stimuli with high disparity contrast tend to have smaller, more vertically oriented regions of least disparity variation around the fovea. These results help explain why the size and shape of optimal spatial integration areas reported in Figs. 5,7, and 9. Pooling signals over spatial integration areas that align with the low variance regions result in reliable disparity estimates.

**Figure 11.**
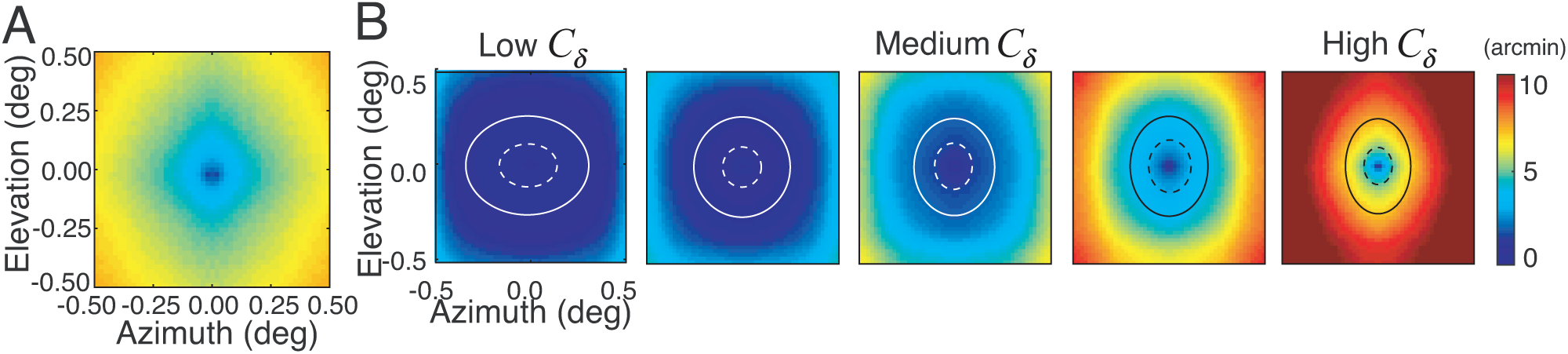
Disparity standard deviation with retinal position and disparity contrast. **A** The standard deviation of natural disparity signals increases systematically with retinal eccentricity. Disparities are more variable at retinal locations farther from foveated points. **B** Same as A, but conditioned on five different disparity contrast bins (left to right [0.1,1,0],[0.2,2.0],[0.4,4.0],[0.75,7.5),[1.5,15] arcmin. At low disparity contrasts, disparities are nearly homogeneous over almost the entire 1° foveal region. At high disparity contrasts, disparity variation increases significantly with eccentricity, and the region of low variability is smaller and vertically elongated. Ellipses (fit by hand) indicate iso-disparity-variation contours.

Figure 12A is identical to Figure 11A, and shows the most likely pattern of disparity variation in our dataset. Outside the central ±1/8°, disparity variance grows linearly with retinal eccentricity, and increases more rapidly with changes in azimuth than with changes in elevation (Fig 12B). Figure 12C-E shows that disparity variance increases less rapidly with eccentricity at far than at near distances (Fig 12C-E). This effect is because the magnitude of a disparity signal for a given depth difference decreases with the square of distance. The effect distance on disparity variation is summarized by the average disparity standard deviation across position as a function of viewing distance (Fig. 12E). Disparity standard deviation decreases monotonically with fixation distance and plateaus at about 15.0m (Fig 12F). Thus, when a far surface is fixated, the disparities in the immediate neighborhood of the fovea are more likely to be near zero. Given that disparity variability decreases with distance, and given that the Burge et al. (2016) dataset contains only object 3m and farther away, it suggests that our estimates of disparity variability are conservative.

**Figure 12:**
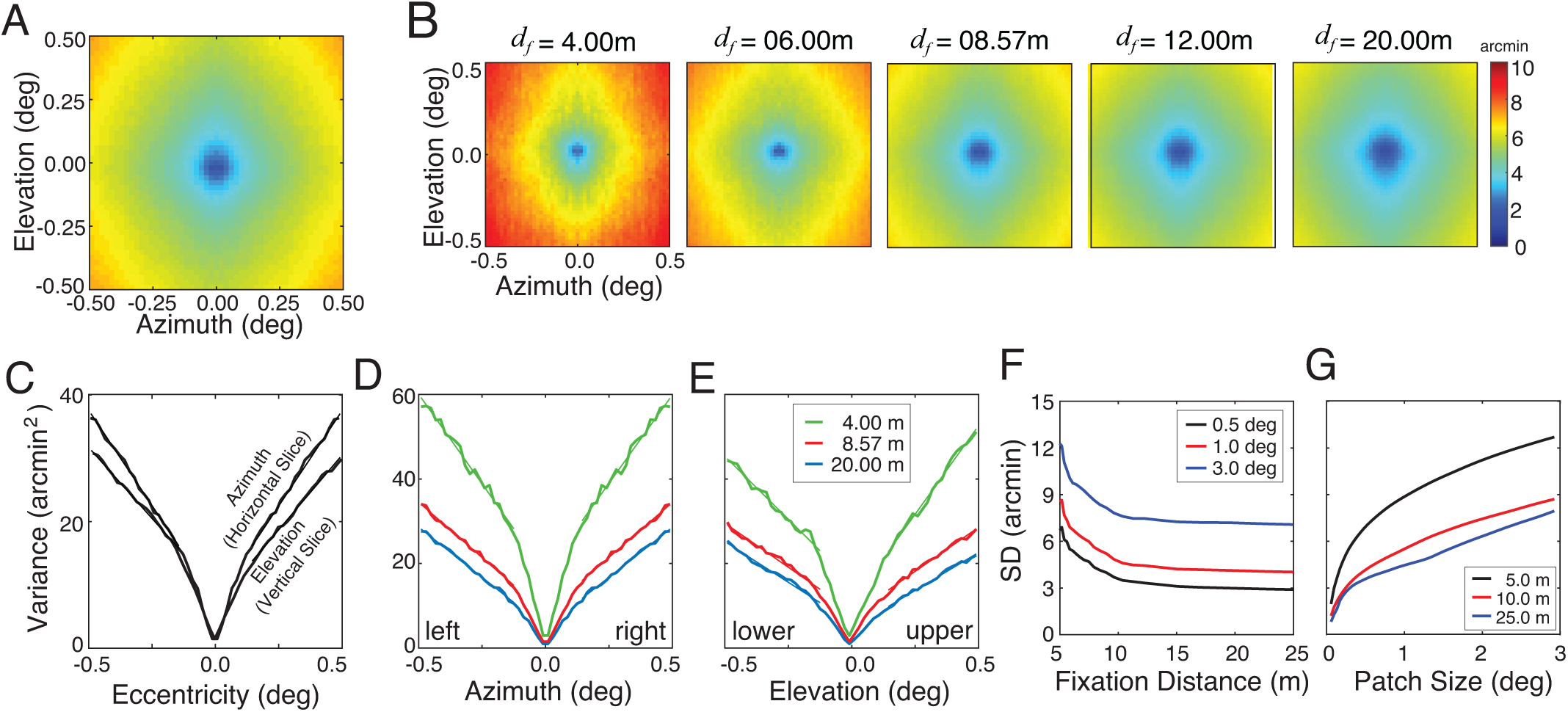
Near-foveal disparities as a function of viewing distance and spatial integration region. **A** Disparity standard deviation across all patches in database (data identical to Fig. 11A). **B** Disparity variance as a function of azimuth and elevation. Disparity variance increases linearly outside the central ±1/8°. Variance increases more rapidly in azimuth 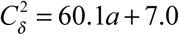 than in elevation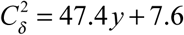 where *a* and *e* are azimuth and elevation in deg, respectively Curves correspond to the squared standard deviation along horizontal and vertical slices through the plot in 11A. **C** Disparity standard deviation at each retinal location, but conditioned on five different viewing distances ranging from 4.0 to 20.0m. For each viewing distance, data is pooled in 0.1 diopter bins centered on the viewing distance. For far distances, disparities near the fovea are more likely to be small. **D,E** Disparity variance in azimuth and elevation as a function of distance (colors). Best fit lines in azimuth range from 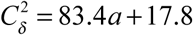 to 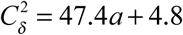 at view distances from 4.0m to 20.0m and best fit lines in elevation range from 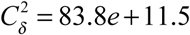 to 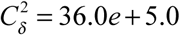 Variance increases more rapidly in the upper than lower visual field. **F** Effect of fixation distance on standard deviation of foveal disparity signals pooled at all eccentricities within a given integration area (colors). Disparity standard deviation decreases monotonically with fixation distance. **G** Effect of patch size on disparity standard deviation for three different distances (colors). Disparity standard deviation increases monotonically with patch size. Distance bins were 0.2 diopters wide.

The spatial area (patch-size) over which the disparities are pooled also impacts the standard deviation (Fig 12G). When the spatial integration area is large, non-negligible surface curvature and depth discontinuities become increasingly likely, and standard deviation increases. When the integration is small, local depth structure is more likely to be well approximated by a locally planar surface (i.e. zero disparity variation). This result is not surprising, but it is important to take note of. How often is local depth structure likely to be well-approximated by a locally flat surface? In our database, the median mean-disparity within a patch is zero and the median standard deviation is less than 0.5 arcmin (see Fig. S3). Nearly planar surfaces are quite common in natural scenes.

Finally, the most likely pattern of disparity variation depends not just on the depth structure of natural scenes but on which scene points are fixated. One weakness of the Burge et al. (2016) dataset, upon which this manuscript is based, is that it has no information about human eye movements. Other datasets do (Gibaldi, Canessa, & Sabatini, 2017; Liu et al., 2008; Sprague et al., 2015). At near distances, humans preferentially fixate objects nearer than random fixations. At far distances (i.e. beyond 3m), human fixations and random fixations are hard to distinguish (Sprague et al., 2015). Thus, the results presented in the current manuscript are likely to be representative of disparity variability for human fixations when objects are farther than 3m, but are likely to underestimate disparity variability across the near and far distances encountered in natural viewing (also see Fig. S1 & Fig. S4).

## DISCUSSION

We developed a high-precision stereo-image sampling procedure, and used it along with a recently published dataset (Burge et al., 2016), to demonstrate how natural depth variation impacts performance, and the size and shape of the mechanisms that optimize performance, in half-occlusion detection and disparity detection with natural stimuli. In the discussion section, we discuss limitations of the work presented here, connections to other topics in the literature, and directions for future work.

### Relationship to previous work

The dataset leveraged in this manuscript has some advantages and some disadvantages compared to other recently published datasets. We compare four recently published datasets, and consider the advantages and disadvantages of each with respect to six factors: i) the presence or absence of eye movements, ii) the presence or absence of groundtruth half-occlusions and groundtruth disparities, iii) the spatial resolution of the images, iv) the range of object distances represented in the dataset, v) the diversity of sampled scenes, and vi) the appropriateness of the dataset for use in psychophysical experiments. Each dataset was collected with a different purpose (or set of purposes) in mind, and each is limited by the technology used to collect the data and by choices made by the researchers.

Sprague et al. affixed human observers with a mobile binocular eye tracker and collected binocular image movies of natural scenes as human observers performed everyday tasks around the University of California, Berkeley (Sprague et al., 2015). The dataset contains objects ranging in distance from 0.5m to Inf. The principal aim in collecting the dataset was to estimate the prior probability distribution of binocular disparities encountered by humans in natural viewing. Absolute disparity depends on the 3D structure of the scene, where the observer is located in the scene, and where the observer gazes in the scene. Collecting stereo-images in concert with matched binocular eye movements is therefore necessary to estimate the distribution of disparities encountered by humans, and the dataset is extremely well suited for this aim. There are two primary disadvantages associated with the dataset. The first disadvantage is that groundtruth disparities and groundtruth occlusions are not known. Disparities were instead estimated from the left- and right-eye images via image-based routines. A second disadvantage is that the stereo-images are low spatial resolution (~9pix/deg). Thus, while this dataset is very well suited for estimating disparity statistics in natural viewing, it is ill suited for examining the accuracy of disparity estimation algorithms, for investigating the impact of local disparity variation on disparity estimation performance, or for obtaining natural stimuli for use in psychophysical experiments.

Gibaldi et al. (2017) tracked binocular eye movements of head-fixed human observers viewing two computer generated 3D scenes from different viewpoints on a stereo-display (Gibaldi et al., 2017). The dataset contains objects ranging in distance from only 0.5 to 1.5m. This paper also had the aim of characterizing disparity statistics in natural viewing. Gibaldi et al.’s computer-generated scenes afford access to groundtruth disparities and groundtruth occlusions. The rendered images had comparatively high spatial resolution (~44pix/deg) and, with appropriate calibration, could be suitable for use in psychophysical experiments. All of these features represent important improvements on the weaknesses of the Sprague et al dataset. The first disadvantage of the Gibaldi et al. dataset is that the eye movements were not collected during observer interaction with the environment; eye movements were instead collected during free viewing of static disparity-specified scenes, presented on a haploscope in a laboratory. A second disadvantage is that the dataset contains only two types of scenes—an office desk and a kitchen table—raising the specter of statistical under-sampling. A third disadvantage is that the images were constructed and rendered in software; although the authors undertook a heroic effort to map natural textures onto high-resolution 3D models of real objects, the possibility remains that the resulting stimuli do not accurately capture all relevant aspects of real scenes. Those caveats aside, this dataset has tremendous potential value, and it provide computer-generated stimuli for both computational and psychophysical studies, especially if it can be expanded.

Adams et al. (2016) collected multiple stereo-image pairs, wide-field (i.e. 360°) laser range scans, and wide-field high-dynamic range images of 76 outdoor scenes near Hampshire, UK (Adams et al., 2016). The dataset contained objects ranging from 1m to Inf. This dataset was collected with the immediate aim of characterizing the statistics of 3D surface orientation as a function of viewing elevation in natural scenes, and the authors developed a sophisticated procedure for estimating local surface orientation from the distance data. The dataset is also very well suited for other applications not relevant to the topic of this manuscript. The stereo-images have very high spatial resolution (~160 pix/deg). One disadvantage of this dataset is that it does not include eye movement data, so the impact of natural eye movements on disparity statistics cannot be estimated. A second disadvantage is that only one range scan was captured per scene. With only one range scan, stereo-parallax precludes precise pixel-wise co-registration of the groundtruth distance data with the left- and right-eye photographic images. Thus, although groundtruth disparity could be computed from the distance data, it is impossible to precisely co-register the stereo-image data with the range data at pixel in both the left- and right-eye images.

Burge et al. (2016) collected 99 stereo-images of natural scenes with laser range scans co-registered to each eye’s photographic image around the University of Texas at Austin campus. The dataset contains objects ranging in distance from 3m to Inf. A robotic gantry aligned the nodal points of the camera and the scanner during data acquisition. As a result, every pixel in each eye’s photographic image contains groundtruth distance data from the corresponding range scan from which groundtruth disparities and groundtruth occlusions can be directly computed. The images in the dataset also have comparatively high spatial resolution (~52 pix/deg). These features of the dataset make it particularly well suited for performing analyses of the impact of local disparity variation on disparity estimation. The first disadvantage of this dataset is that it does not contain eye movement data, although the technique used by Gibaldi et al. (2017) could be applied to get comparable data (also see (Liu, Cormack, & Bovik, 2010)). However, because the data has high-spatial resolution and co-registered groundtruth distance information, the dataset should prove useful as a source for psychophysics stimuli and for future computational studies.

### Adaptive filtering in psychophysics and neuroscience

The computational results reported here predict that human performance in disparity-related tasks can benefit from adapting the size and shape of receptive fields to the disparity contrast of each stimulus. Is adaptive filtering based on properties of visual stimuli neurophysiologically plausible? Yes. Increases in luminance contrast are associated with decreases in the spatial size of receptive fields in macaque V1 (Cavanaugh, Bair, & Movshon, 2002; Sceniak, Ringach, Hawken, & Shapley, 1999). Increases in luminance contrast are also associated with decreases in the temporal integration period in macaque V1 and MT (Bair & Movshon, 2004).

### The influence of priors in perception

In recent years, prior probability distributions have received considerable attention. The impact of priors on perceptual biases (Burge et al., 2010; Burge, Peterson, & Palmer, 2005; Girshick, Landy, & Simoncelli, 2011; Parise, Knorre, & Ernst, 2014; Stocker & Simoncelli, 2006; Weiss, Simoncelli, & Adelson, 2002) and on the design of neural systems (Liu et al., 2008; Sprague et al., 2015) have been extensively investigated. However, Bayesian estimation theory predicts that priors should significantly impact perceptual estimates only when measurements are highly unreliable (Knill & Richards, 1996). Factors other than the prior are likely to be more important determinants of performance and system design than the prior in many (most?) viewing situations (Burge & Jaini, 2017).

Psychophysics is principally concerned with understanding the lawful relationships between stimulus properties and human performance in critical tasks. Human performance in natural tasks varies from stimulus to stimulus because stimuli differ in their task-relevant properties. The prior probability distribution cannot account for stimulus-to-stimulus performance variation. Thus, it is necessary to characterize stimulus variability and develop models that predict its impact on psychophysical performance. A great deal of previous work has examined the impact of external noise on performance (Geisler & Davila, 1985; Pelli, 1985). Comparatively little work has systematically examined the impact of natural stimulus variability on performance (but see (Burge & Geisler, 2011; 2014; 2015; Geisler & Perry, 2009; Hibbard, 2008)). The current paper takes an important step towards the aim of predicting in the context of two tasks related to disparity-processing.

### Stereo-image patch sampling for psychophysics

Task-specific computational analyses, like those presented here, are useful for determining the optimal solutions to sensory-perceptual problems, and for developing targeted hypotheses about biological visual systems. To determine whether the results presented are in fact relevant to biological visual systems, psychophysical experiments are ultimately required. The stereo-image sampling and interpolation procedure developed here should facilitate future experiments on human disparity processing and stereopsis with natural stimuli.

## CONCLUSION

In this manuscript, we developed a high-fidelity stereo-image sampling procedure and used it to investigate the impact of local depth variation on half-occlusion detection and disparity detection in natural scenes. Local depth variation decreases the size and vertically elongates the shape of the binocular spatial integration area that optimizes performance.

## METHODS

### Contrast images and binocular difference images

The inputs to the human visual system are the left- and right-eye retinal images. Disparity-processing mechanisms are widely modeled to operate on local contrast signals, the output of luminance normalization mechanisms in the retina. The Weber contrast image ***c*** is obtained from a luminance images *I* by subtracting off and dividing by the mean

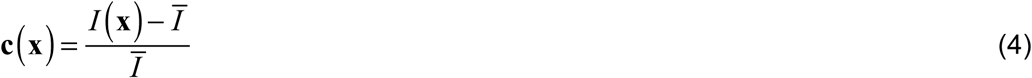

Where 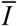 is the local windowed mean and **x**_0_ = (*x*_0_, *y*_0_) is the location of the central pixel. The local windowed mean is given by

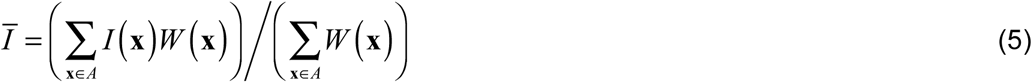

where *W* (**x**) is a spatial windowing function. We have used Gaussian or raised-cosine windowing functions; results are highly robust to the specific type of window. The windowed Weber contrast image

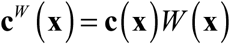

is obtained by point-wise multiplying the Weber contrast image by the window.

### Binocular difference image contrast

The binocular difference image is given by the point-wise difference of the two retinal images

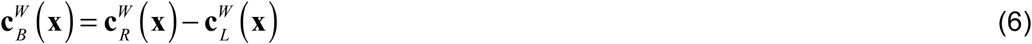

where ***c***_*L*_ and ***c***_*R*_ are the windowed left- and right-eye contrast images where the center pixels of each image are centered on candidate corresponding points (see Fig. 2A,B). Thus, the binocular difference image is the point-wise difference of the left-and right-eye contrast images. The RMS contrast of the binocular difference image *C*_*B*_ is given by

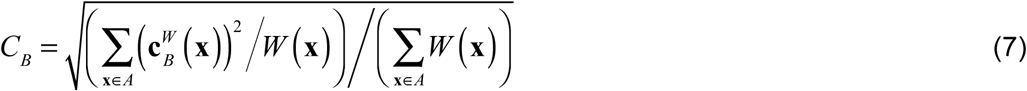

where *W* (**x**) is the window that imposes the spatial integration area.

### Binocular disparity contrast

Our sampling procedure ensures that the center of each sampled stereo-image patch corresponds to the same surface point in the scene, assuming that the surface point is binocularly visible. If the surface point is half-occluded (i.e. visible to only one eye), its image only falls on the fovea of the anchor eye and a point on the occluding surface will be imaged at the fovea of the other eye. Disparity is undefined at half-occluded points, so we compute disparity contrast only for binocularly visible points.

To compute disparity, a point of reference must be assumed. We compute disparity relative to the center pixel of the anchor eye’s image patch (see Results). This computation is equivalent to computing absolute disparity, assuming that the center pixel of the anchor eye’s image corresponds to a binocularly visible scene point and that the eyes are fixating it. It is also equivalent to computing relative disparity where the point of reference is the center pixel of the anchor eye’s image. To compute the groundtruth disparity pattern from groundtruth distance, we first compute the vergence demand at each pixel from the distance data, and then subtract the vergence demand at the central pixel from the vergence demand of every other pixel in the patch. All vergence angles are computed in the epipolar plane. The result is the pattern of absolute near-foveal disparities *δ* (**x**) that would result from fixating the surface point in the scene corresponding to the center pixel of the anchor eye’s image.

Root-mean-squared (RMS) disparity contrast is a scalar measure of variation about the mean in a local spatial area. The RMS disparity contrast is given by

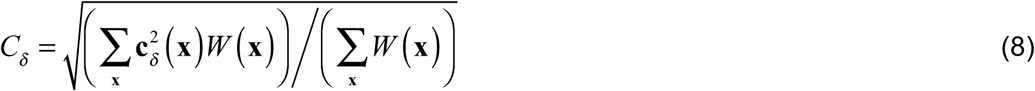

where 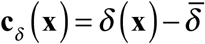 is the mean-centered disparity image.

### Comparing detection sensitivities for fixed and adaptive spatial integration areas

In a two-presentation forced choice task, proportion correct *P* is given by the area under the ROC curve (e.g. Fig. 5D). The corresponding sensitivity *d* ' is given by

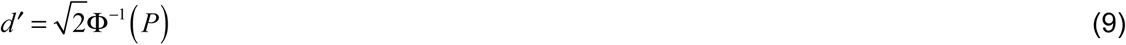

where Ф^-1^ (.) is the inverse cumulative normal. With fixed spatial integration areas, window size is fixed to maximize sensitivity across all stimuli regardless of disparity contrast. With adaptive filtering, window size changes to optimize sensitivity at each disparity contrast. Overall proportion correct with adaptive filtering is given by a weighted sum of the proportion correct *P*_*i*_ in each non-overlapping disparity-contrast bin

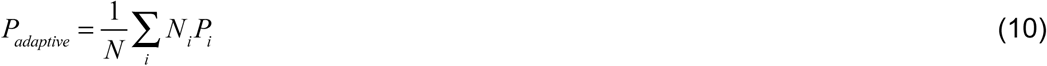

where *N*_*i*_ is the number of stimuli in disparity-contrast bin *i*.

## Supplement The effect of depth variation on disparity tasks in natural scenes

**Figure S1.**
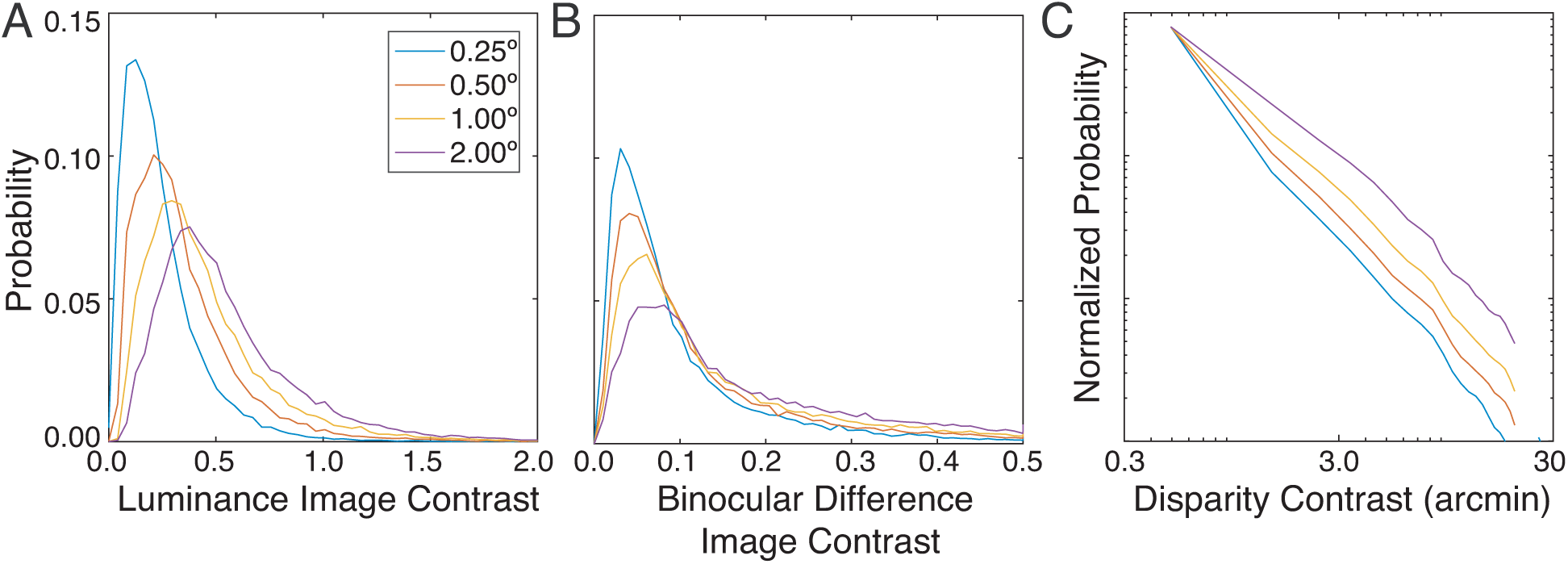
Luminance contrast, binocular difference image contrast, and disparity contrast in natural scenes. **A** Distribution of luminance contrast for different spatial integration areas (colors). Larger luminance contrasts are more likely when computed with larger spatial integration areas (see Methods) **B** Distribution of binocular difference image contrast. Binocular difference image contrast is the contrast of the point-wise difference between the right- and left-eye windowed contrast images (see Methods). Low binocular difference image contrasts are associated with low disparity contrasts (see Fig. 10). **C** Distribution of disparity contrasts. Disparity contrast is approximately distributed as a power law, although the approximation breaks down at very small and large contrasts. The y-axis spans three-orders of magnitude. Note that because the database of natural scenes from which these statistics are calculated contain no objects closer than 3.0m, and because the disparity associated with a given depth decreases with the square of distance, these distributions probably underestimate the frequency of high disparity contrasts.

**Figure S2:**
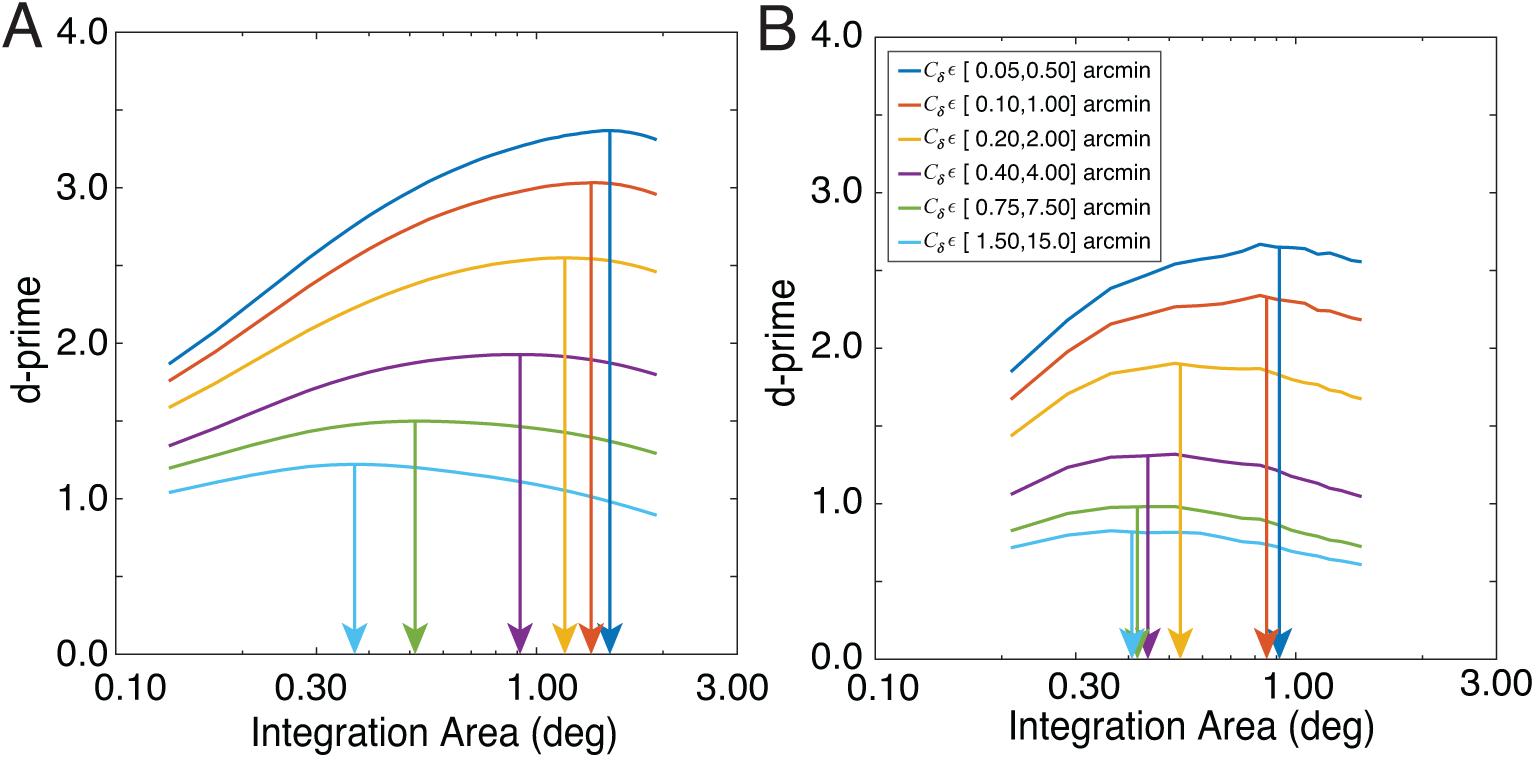
Effect of disparity contrast on task-performance and size of task-optimal integration region **A** Half-occlusion sensitivity as a function of integration area for different disparity contrast intervals (colors). Arrows mark the spatial integration area at half-height for which half-occlusion detection performance is optimized. The analyses are the same as in Fig 5E but for a different choice of disparity contrast bins in arcmin. The dashed curve shows the d-prime obtained for the superset of stimuli in all disparity contrast intervals. **B** Disparity-detection sensitivity as a function of integration area for different disparity contrast intervals (colors). Similar to **A**, but for the disparity detection task (see also Fig 7E)

**Fig S3:**
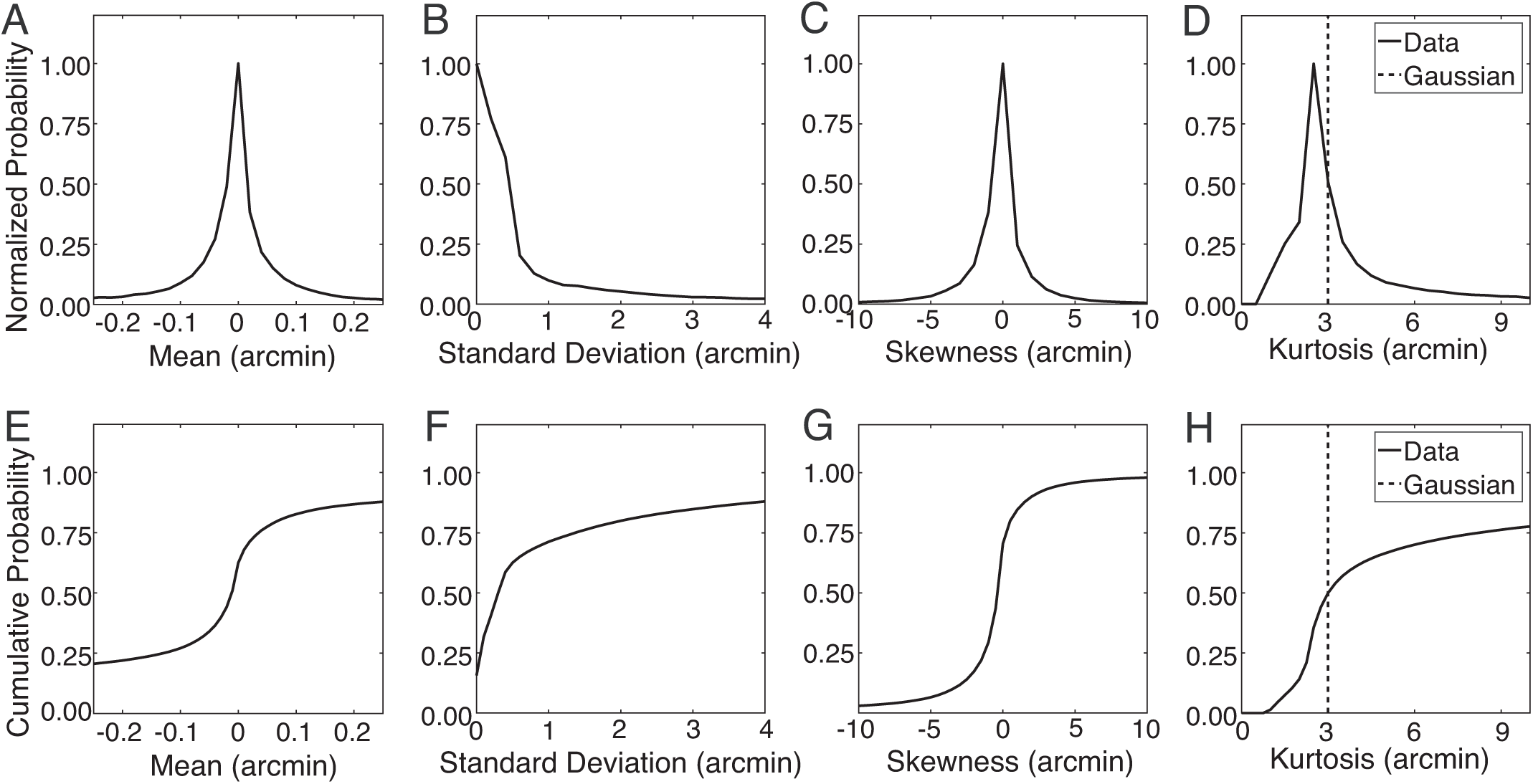
Distributions of within-patch statistics. **A** The distribution of within-patch mean disparity, **B** standard deviation, **C** skewness, and **D** kurtosis, aggregated across all patches in the database. The most likely median and skewness are both zero, suggesting that the within-patch distribution is frequently symmetric about zero. The median skewness is near 0.0 and the median kurtosis is near 3.0, suggesting that within each patch, the median distribution of within-patch disparities is approximately Gaussian. This result suggests that the median within-patch distribution of disparities is well-approximated as Gaussian. However, for a given distribution to be approximately Gaussian, skewness and kurtosis need to take on the appropriate values within the same patch. To check, we computed the joint histogram of within-patch skewness and kurtosis. Results confirm that the median natural image patch is a nearly planar surface with Gaussian distributed disparities.

**Figure S4:**
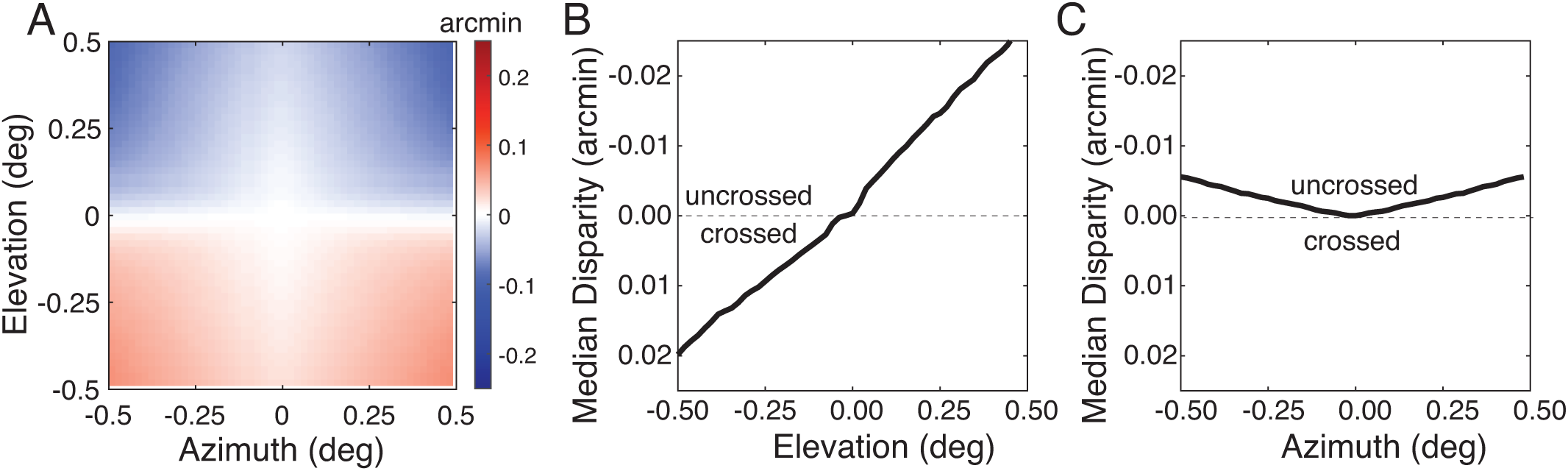
Median disparities in the near foveal region. **A** The spatial map of natural disparity medians is symmetric about the vertical meridian. Uncrossed disparities occur exclusively at elevations above the horizontal median. Crossed disparities occur exclusively at elevations below the horizontal meridian. This pattern of disparities can be understood in terms of the typical structure of natural scenes where nearer fixations (yielding crossed disparities) occur most frequently at lower elevations where the ground-plane dominates. **B** Vertical horopter. A vertical slice in A at an azimuth of 0.0° shows a backward pitch with respect to the vertical, consistent with a previously reported property of the empirical vertical horopter known as the Helmholtz shear. The best-fit line is *δ* _*median*_ = -0.0475*e* - 0.0028. The top-back pitch of the vertical horopter is consistent with multiple psychophysical observations. **C** Horizontal horopter. Median disparities along the horizontal median are more uncrossed at larger azimuths. A horizontal slice in A at an elevation of 0.0° is well fit by the parabola *δ median* = - 0.022*a*^2^. The sign of the horopter’s inflection (toward uncrossed disparity) is consistent with the Hering-Hillebrand deviation.

It has been proposed that retinal corresponding points are positioned to be most often stimulated by surfaces in the world (Helmholtz, 1925). This proposal can account for why the empirical vertical horopter is pitched back (i.e. the Helmholtz shear) and why the empirical horizontal horopter deviates from the Vieth-Müller circle (i.e. the Hering-Hillebrand deviation). Median disparities along the vertical and horizontal retinal meridians constitute predictions of the empirical horizontal and vertical horopters. We computed the median disparities along the horizontal and vertical retinal meridians in our dataset (Fig. S4A-C). The median disparities exhibit the Helmholtz shear and the Hering-Hillebrand deviation, but smaller in magnitude from psychophysically reported values, consistent with the expectation that disparity reliability is underestimated because of the absence of objects nearer than 3m in our dataset.

